# CONSTRUCT: an algorithmic tool for identifying functional or structurally important regions in protein tertiary structure

**DOI:** 10.1101/2024.10.07.617015

**Authors:** Lucas Chivot, Antoine Bridier-Nahmias, Loïc Favennec, Jean-Christophe Gelly, Jérôme Clain, Romain Coppée

**Author notes:** Corresponding author: Romain Coppée Laboratoire de Parasitologie-Mycologie Unité de Recherche 7510 ESCAPE Université de Rouen Normandie Rouen, F-76000 France.

## Abstract

In the realm of protein-coding genes, evolutionary rates show considerable variability. Essential or highly expressed proteins evolve more slowly, and within a protein, different amino acid sites evolve at different rates. Accurately modeling this variation is critical for identifying structurally or functionally important amino acid sites. Standard methods such as Rate4Site assume independent substitution rates across sites, and the most conserved ones are widely distributed in protein tertiary structure. This is biologically unrealistic because functional sites often cluster together. Here, we developed CONSTRUCT, an improved strategy for identifying functional and structurally important regions in protein tertiary structure. Given a set of orthologous sequences, CONSTRUCT first calculates site-specific substitution rates using the Rate4site model, which are then weighted by the rates of neighboring amino acid sites within a range of window sizes. The optimal window size is determined by the strongest spatial correlation, if present. This method can be analyzed using either Cα atoms or the center of mass of amino acid sites to account for side chain orientation. Validation on 14 functionally characterized proteins of different size, conservation level and kingdom/domain of species has demonstrated the relevance of CONSTRUCT.

## Introduction

In the realm of protein-coding genes, evolutionary rates vary widely within a species. Genes encoding essential or abundantly expressed proteins evolve more slowly (1, 2). Within the same protein, amino acid sites also exhibit different rates of evolution. Some of this variation in evolutionary rates arises from selection, either positive such as adaptation to environmental changes or negative (*id est* purifying selection) to maintain critical protein activities (3). Accurately modeling this variation is critical in evolutionary studies (4), particularly for identifying amino acid sites within proteins that are structurally or functionally important (5).

Over the years, research has shown that site-specific evolutionary rates are shaped by both structural and functional constraints (6–8). In particular, less critical parts of a molecule tend to evolve more rapidly than essential ones (2). Advances in computational evolutionary modeling have yielded robust methods for estimating site-specific substitution rates from amino acid and DNA sequences (3, 4, 9, 10). Comparison of orthologous sequences (gene sequences in different species that have evolved from a common ancestor through speciation) reveals conserved amino acid sites that are likely to be structurally and/or functionally important (5). These amino acid sites tend to evolve under intense purifying selection to maintain protein function.

Several bioinformatics tools based on phylogenetics have been developed to identify functional amino acid sites (5, 11–14). These tools have shown that conserved sites can align with experimental evidence, validating their utility despite several limitations. Widely used tools such as Rate4Site assume that site-specific substitution rates are independent across amino acid sites and identically distributed in the sequence alignment (14). This assumption simplifies statistical modeling, but is biologically unrealistic. It implies that slowly evolving functional sites are randomly scattered throughout the protein tertiary structure. In reality, functionally important amino acid sites often cluster to form regions such as binding or catalytically active sites (15, 16). Thus, this assumption is inappropriate for modeling functional regions composed of multiple amino acid sites.

To address this limitation, some methods have been introduced to account for the spatial correlation of evolutionary patterns. Many of these methods employ a sliding window framework that approximates the site-specific substitution rate for a given amino acid site by the average of neighboring sites within the protein tertiary structure (17–19). An amino acid site is considered a neighbor if its Euclidean distance from the focal site is within a predefined window size. However, sliding window methods have inherent disadvantages. First, most methods give equal weight to the focal site and its neighbors, even though the focal site contains more pertinent information about its substitution rate. Second, determining the optimal window size is challenging. A window that is too large will include too many distant sites, potentially biasing the estimate of the target amino acid site, while a window that is too small will fail to capture spatial correlation and may lead to overfitting. In addition, there is evidence that the optimal window size may vary between different protein families (17).

A novel phylogenetic Gaussian process model has been developed to identify conserved amino acid patches in protein tertiary structure (20). This approach computes the best characteristic length for estimating spatially correlated site-specific substitution rates, thus capturing functional regions. However, the model implemented in the GP4Rate software has drawbacks. It relies on the Markov chain Monte Carlo method to generate samples, which can be time-consuming (from hours to days, depending on the size of the dataset). This limitation has been addressed by the FuncPatch web server, which is designed as a fast approximation to GP4Rate (21). In addition, the GP4Rate method searches for spatial correlation of site-specific substitution rates based on Cα atoms, ignoring the orientation of amino acid side chains. However, including side chain orientation is critical for helix-enriched proteins such as transmembrane proteins. Finally, despite the relevance of GP4Rate and FuncPatch, these tools are no longer available.

In this study, we developed an improved strategy called CONSTRUCT to identify patches of conserved sites within protein tertiary structures as putative functional and/or structurally important regions. Site-specific substitution rates are first calculated using the Rate4Site model and then weighted by the site-specific substitution rates of neighboring amino acid sites according to Euclidean distances, over a range of window sizes (from 1 to 20 Ångströms (Å)). The best window size is then chosen by selecting the strongest spatial correlation of site-specific substitution rates, if present. The analysis can be performed either on Cα atoms or using the center of mass of amino acid sites as a *proxy* for side chain orientation. To demonstrate the accuracy of CONSTRUCT in identifying critical protein regions, 14 case studies were performed on various functionally characterized proteins (transport and membrane proteins, enzymes, oncoproteins) of different sizes, levels of interspecies conservation, and kingdoms or domains (*Metazoa*, *Fungi*, *Chromalveolata*, *Bacteria*).

## Materials and Methods

### Calculation of site-specific substitution rates

The calculation of site-specific evolutionary rates was performed using the Rate4Site computational tool (14) with an amino acid sequence alignment as input (**Figure 1A**). The user must ensure the quality of the alignment and remove gapped and misaligned regions. Rate4Site infers a phylogenetic tree to model the evolutionary relationships among the sequences, which is used to weight the calculation of site-specific substitution rates. Rate4Site assumes that the provided multiple sequence alignment represents orthologous sequences and that each site in the alignment corresponds to the same position in all sequences. The algorithm is implemented in a Bayesian framework and uses a random-effects approach, specifying gamma distributions as the prior rate distribution. An Empirical Bayes approach is then used to calculate site-specific substitution rates. The output is a score for each amino acid site, where lower scores indicate more conserved amino acid sites (likely under purifying selection) and higher scores indicate more variable amino acid sites.

**Figure 1.**
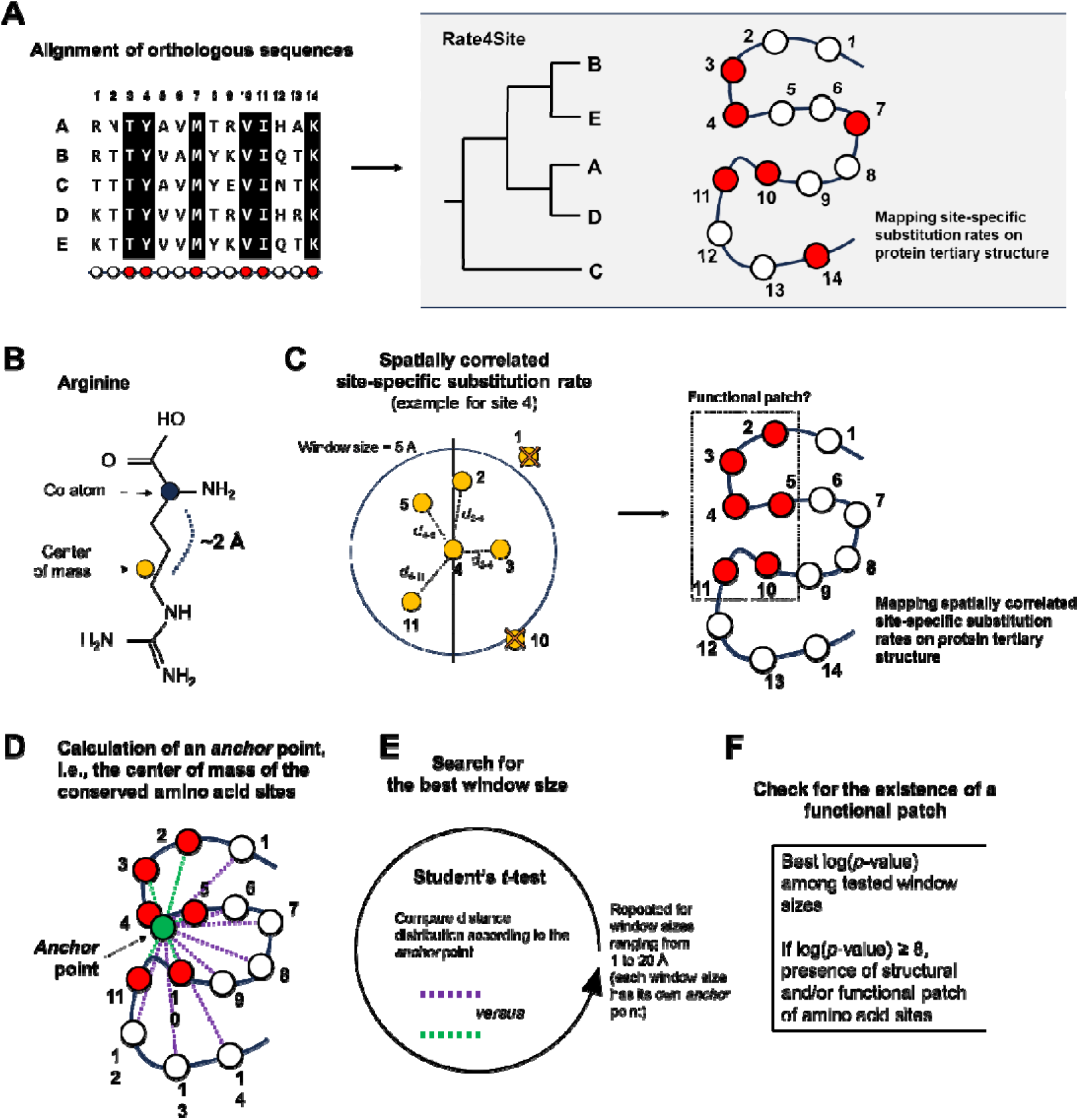
- Overview of the CONSTRUCT algorithm. **(A)** CONSTRUCT requires a multiple orthologous sequence alignment and a representative tertiary structure. A phylogenetic tree is constructed to infer site-specific substitution rates using Rate4Site, which ignores the spatial correlation of site-specific substitution rates in protein tertiary structure. **(B)** The Euclidean distance between amino acid sites is calculated using either the C_α_ atoms or the center of mass of amino acids as a *proxy* for side chain orientation (here using an arginine as an example). **(C)** For an amino acid site, its site-specific substitution rate is weighted by the rates of its neighboring residues; this weighting is distance-dependent: the influence of a neighbor’s substitution rate decreases as the Euclidean distance between residues increases. In this example, the spatially correlated site-specific substitution rate is calculated for site 4 in a window size of 5 Å (a range of distances from 1 to 20 Å is performed with a step size of 1 Å). The spatially correlated site-specific substitution rate is calculated for all sites, then the most conserved sites are highlighted on the protein tertiary structure. **(D)** One *anchor* point per protein is calculated as the geometric center of the most conserved region of the protein. **(E)** A Student’s *t*-test is performed to compare the Euclidean distances of the top 10% conserved amino acid sites to the *anchor* point with the Euclidean distances of the remaining amino acid sites to the *anchor* point. The analysis is performed for each window size (one anchor point is calculated for each window size). **(F)** A log(*p*-value) greater than or equal to 8 indicates the presence of a spatial correlation in site-specific substitution rates. Only the results with the window size that maximizes the spatial correlation are kept.

### Estimation of spatially correlated site-specific substitution rates

Once the site-specific substitution rates have been estimated, these rates are further refined by incorporating the structural context of the protein. This process requires the user to provide a tertiary structure of the protein in PDB format. The reference sequence from the multiple sequence alignment and the sequence extracted from the PDB file are aligned to map the site-specific substitution rates to the corresponding amino acid sites in the tertiary structure.

After mapping the substitution rates to the structure, a predefined window size is used to identify neighboring amino acid sites for each residue in the protein. Two approaches can be used to define these neighbors: one based on the peptide backbone, specifically Cα atoms, and the other based on the center of mass of each amino acid site (**Figure 1B**). The center of mass approach takes into account the spatial orientation of the amino acid side chains, providing a likely more accurate representation of the local structural environment.

For each amino acid site, its site-specific substitution rate is then weighted by the rates of its neighboring residues (**Figure 1C**). This weighting is distance-dependent, meaning that the influence of a neighbor’s substitution rate decreases as the Euclidean distance (expressed in Å) between the residues increases.

### Detection of conserved amino acid patches

First, conserved amino acid patch(es) can be identified by eye by looking at the spatially correlated site-specific substitution rates mapped on the protein surface. Second, to investigate formally the existence of a conserved amino acid patch, we implemented a statistical approach. An artificial anchor point is generated within the three-dimensional structure of the protein. This anchor point is defined as the center of mass of the top 10% of amino acid sites with the lowest spatially correlated site-specific substitution rates (i.e., the geometric center of the most conserved amino acid sites of the protein) (**Figure 1D**). The rationale behind selecting the top 10% of most conserved amino acid sites is to focus on those residues that are likely to have critical functional or structural roles due to their high conservation. Next, the Euclidean distance (in Å) between the anchor point and each amino acid site (either the Cα atoms or the center of masses, depending on the user choice) is calculated. These distances provide a measure of how spatially close is each amino acid site to the most conserved region of the protein.

The Euclidean distances to the anchor point of the top 10% conserved amino acid sites are then compared to those of the remaining amino acid sites by a Student’s *t*-test (**Figure 1E**). A significant difference between these two sets of distances indicates a spatial clustering of conserved residues around the anchor point.

After performing the statistical test, the logarithm of the *p*-value obtained from the test is calculated. This log-transformed *p*-value provides a more interpretable measure of statistical significance. A log(*p*-value) greater than or equal to 8 serves as a conservative threshold to confirm the existence of a spatial correlation in site-specific substitution rates (**Figure 1F**). It also confirms the presence of a conserved amino acid patch that is likely to be critical for the function and/or structural integrity of the protein.

### Determining the best window size for spatially correlated site-specific substitution rates

The strength of spatial correlation of site-specific substitution rates can vary among proteins and also depends on the window size chosen. Therefore, the calculation of site-specific substitution rates weighted by the three-dimensional environment is performed over a range of distances from 1 to 20 Å, with a step size of 1 Å (**Figure 1E**). The upper limit of 20 Å is based on the rationale that an amino acid site that is excessively distant from the residue under study may have a negligible effect on its evolutionary rate.

For each window size within this range, spatially correlated site-specific substitution rates are calculated. This results in 20 different sets of spatially correlated site-specific substitution rates. Distance comparisons are made between the top 10% conserved amino acid sites or the less conserved amino acid sites and the anchor point, independently for each window size. The 20 log-transformed *p*-values are then calculated and compared. Among the 20 window sizes, the highest log(*p*-value) is considered the optimal value, as it represents the system that maximizes the strength of the spatial correlation. This approach ensures that the selected window size best captures the local structural influences on the evolutionary rates.

### Application of CONSTRUCT to case studies

All of the tasks described above have been integrated into a software package called CONSTRUCT. To demonstrate the accuracy of CONSTRUCT in identifying functional amino acid patches, we developed a series of datasets for several functionally characterized proteins, including transport and membrane proteins, enzymes, and oncoproteins. These proteins vary in size, conservation levels across species, and originate from different kingdoms/domains including mammals, parasites, and bacteria (**Table 1**). Here we have focused on cytochrome c, E3 ubiquitin-protein ligase MDM2, GTPase HRas, myoglobin, cystic fibrosis transmembrane conductance regulator (CFTR), cAMP-dependent protein kinase catalytic subunit alpha, mitogen-activated protein kinase 1, sodium/glucose cotransporter 1 (SLGT1), Kelch-like ECH-associated protein 1 (KEAP1), and torsin-1B in mammals; the dihydropteroate synthase (DHPS) and dihydrofolate reductase (DHFR) enzymes in parasites; the aromatic amino acid exporter YddG in bacteria; and the GDP-mannose transporter 1 in fungi. For each protein, detailed information was compiled, including its function, cellular localization, conserved domains, amino acid sites reported to be functionally important, and a description of its three-dimensional structure (**Supplementary Data S1**). In one case, the three-dimensional structure was not resolved and was predicted using the AlphaFold 3 algorithm (DeepMind, Google) (22) to show that CONSTRUCT can also identify relevant functional regions with predicted structures. All results were compared with those of Rate4Site, which ignores the spatial correlation of site-specific substitution rates in protein tertiary structure (14). Unless otherwise specified, the calculation of spatially correlated site-specific substitution rates was based on the center of mass of amino acid sites as Cartesian coordinates. The execution time of the software was benchmarked across the case studies on a Linux system (Ubuntu 22.04.4 LTS) consisting of 16 Go random access memory (RAM) and an Intel Core I7-6820HQ (**Supplementary Table S1**).

**Table 1.** – Characteristics of the proteins investigated in this study.

| Protein | Classification | Main function | Kingdom or class | Reference species | Uniprot ID | Protein Length | Protein domain investigated | PDB structure | # orthologous sequences | Alignment length | Conservation level among orthologous (%) |
| --- | --- | --- | --- | --- | --- | --- | --- | --- | --- | --- | --- |
| Cytochrome c | Transport protein | Electron transfer | Metazoa | *Homo sapiens* | P99999 | 105 | — | 1J3S | 853 | 103 | 85.15 |
| MDM2 | Oncoprotein | Regulates p53 | Metazoa | *Homo sapiens* | Q00987 | 491 | N-terminal domain (25-109) | 1YCR | 376 | 109 | 92.10 |
| KEAP1 | Peptide binding protein | Regulates Nrf2 | Metazoa | *Homo sapiens* | Q14145 | 624 | Kelch domain (325-609) | 2FLU | 135 | 285 | 87.89 |
| Myoglobin | Transport protein | Storage and transport of oxygen | Metazoa | *Homo sapiens* | P02144 | 154 | — | 3RGK | 454 | 153 | 86.83 |
| CFTR | Membrane transport protein | Chloride ion transport, regulation of salt and water transport | Metazoa | *Homo sapiens* | P13569 | 1,480 | — | 8FZQ | 415 | 1463 | 88.47 |
| cAMP-dependent protein kinase A | Transferase | Phosphorylation of serine and threonine on target proteins | Metazoa | *Homo sapiens* | P17612 | 351 | — | 4WB5 | 254 | 334 | 97.53 |
| DHFR | Oxidoreductase | Catalyzes the reduction of dihydrofolate to tetrahydrofolate | Chrom-alveolata | *Plasmodium falciparum* | A7UD81 | 608 | N-terminal domain (1-231) | 3QGT | 366 | 617 | 48.53 |
| DHPS | Transferase | Catalyzes the formation of dihydropteroate | Chrom-alveolata | *Plasmodium falciparum* | Q25704 | 706 | Catalytic domain (383-708) | 6JWV | 55 | 641 | 64.15 |
| Mitogen-activated protein kinase 1 | Transferase | Transmits extracellular signals to the nucleus | Metazoa | *Rattus Norvegicus* | P63086 | 358 | — | 5UMO | 493 | 365 | 73.93 |
| SGLT1 | Membrane transport protein | Glucose and galactose transport | Metazoa | *Homo sapiens* | P13866 | 664 | — | 7SL8 | 402 | 667 | 79.34 |
| Torsin-1B | Chaperone protein | Participates in protein folding | Metazoa | *Homo sapiens* | O14657 | 336 | — | Predicted | 299 | 338 | 81.08 |
| YddG | Membrane transport protein | Transport of aromatic amino acids | Bacteria | *Ancylobacter novellus* | D7A5Q8 | 287 | — | 5I20 | 383 | 277 | 63.47 |
| GDP-mannose transporter 1 | Membrane transport protein | GDP-mannose transport | Fungi | *Saccharomyces cerevisiae* | P40107 | 337 | — | 5OGE | 366 | 322 | 66.53 |
| HRas | Oncoprotein | Ras protein signal transduction | Metazoa | *Homo sapiens* | P01112 | 189 | — | 5P21 | 421 | 166 | 93.62 |

### Implementation

CONSTRUCT can be run on any computer with a Linux system and the Python 3 and R programming languages installed. The software can be easily installed using a bash script that checks for dependencies (**Supplementary Table S2**). To simplify the use of CONSTRUCT, a graphical user interface has been developed. The CONSTRUCT program and documentation are freely available on github (https://github.com/Rcoppee/CONSTRUCT). A video tutorial has been created for the installation and use of CONSTRUCT: https://youtu.be/bf-VYReZIeM.

## Results

### Defining the cutoff to detect patch of conserved amino acid sites

In CONSTRUCT, we set a log(*p*-value) ≥ 8 as a conservative threshold to confirm the existence of a spatial correlation in site-specific substitution rates. To establish this threshold, we calculated the log(*p*-value) for proteins of different sizes after performing random permutations of the site-specific substitution rates. These permutations destroy the spatial correlation of the site-specific substitution rates. Following the permutations, the CONSTRUCT algorithm was run to compute spatially correlated site-specific substitution rates in the range of 1 to 20 Å. This random permutation was applied to 1, 000 independent replicates, generating a total of 20, 000 log(*p*-value). In parallel, the analysis without permutation was performed to verify the presence of functional amino acid patches, which were then cross-referenced with functional data from the literature. This strategy was applied to the MDM2 protein (a small protein of around 80 amino acids that was also studied in the Huang and Golding paper to illustrate FuncPatch) (23) and the cAMP protein (a protein of about 350 amino acids) (24). For that, two datasets of 376 and 254 orthologous sequences were generated for MDM2 and cAMP, respectively (**Table 1**).

Using only the Rate4Site algorithm, which ignores the spatial correlation of site-specific substitution rates in protein tertiary structure, conserved amino acid sites were distributed throughout the protein structure in both MDM2 and cAMP (**Figure 2A** and **2D**). Using CONSTRUCT and in the absence of permutation, a spatial correlation of site-specific substitution rates was detected for both MDM2 and cAMP. For MDM2, the maximum strength of spatial correlation was observed at a distance of 10 Å, associated with a log(*p*-value) of 15.07 (**Table 2**). The most conserved sites identified by CONSTRUCT formed a well-defined patch in the tertiary structure (**Figure 2B**). In addition, the conserved patch of amino acid sites overlapped with the MDM2 cleft, which directly interacts with the p53 protein (23, 25). Therefore, the predicted conserved patch is likely to be important for the MDM2-p53 interaction. Similarly, for cAMP, the maximum strength of spatial correlation was detected at a distance of 17 Å, associated with a log(*p*-value) of 78.87 (**Table 2**). Again, the most conserved sites identified by CONSTRUCT formed a well-defined patch in the tertiary structure previously reported as the site of ATP interaction (**Figure 2E**). In particular, the patch included the amino acid sites Arg165, Asp166, Lys168, Thr195, Tyr204, Leu205, Glu208, Tyr215, Asp220, Trp222, and Arg280, which are important for catalytic activity (24, 26).

**Figure 2.**
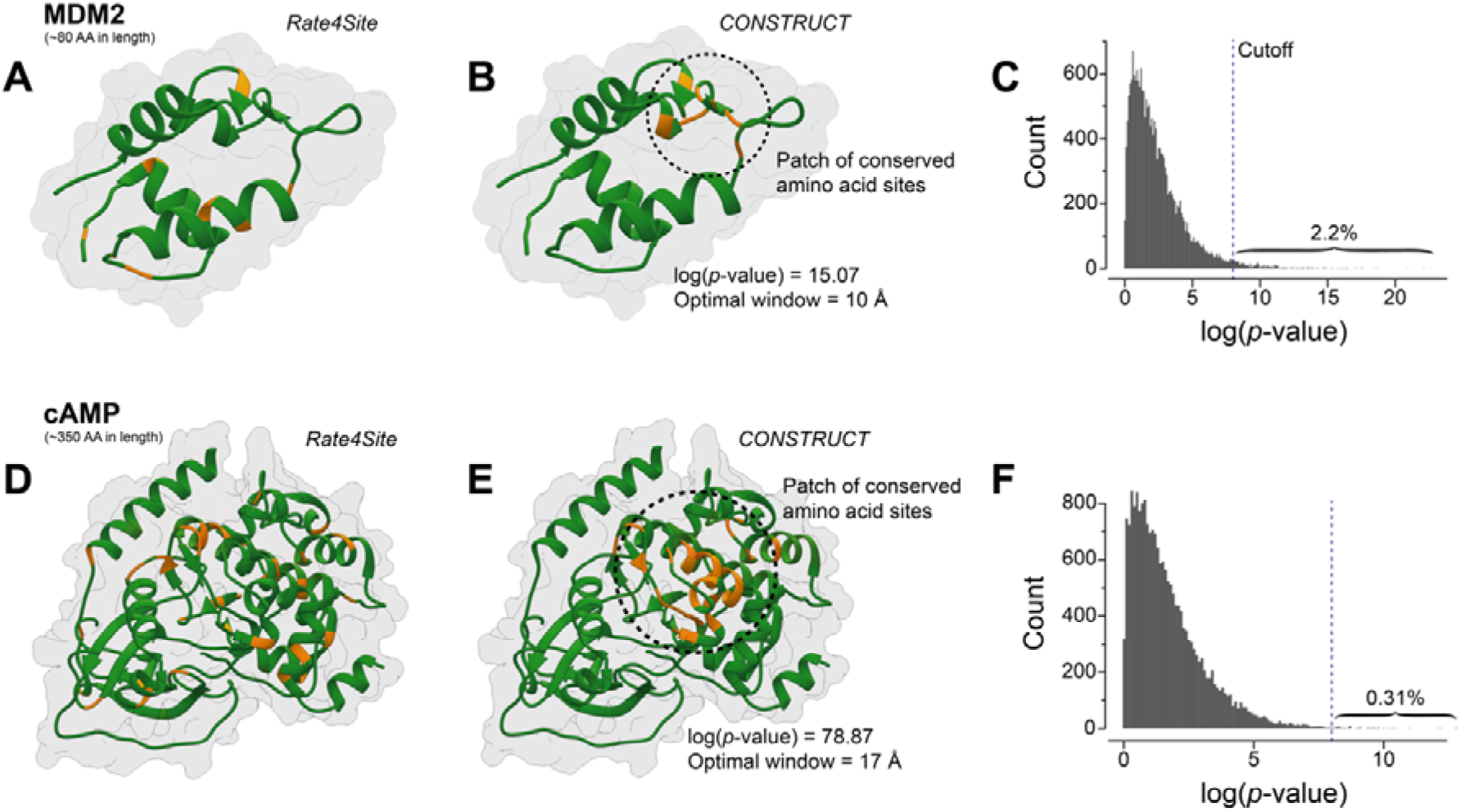
– Defining the cutoff to detect patches of conserved amino acid sites. Location of the 10% most conserved amino acid sites (colored in orange) in the tertiary structure of MDM2 and cAMP according to **(A** and **D)** Rate4Site and **(B** and **E)** CONSTRUCT. **(C** and **F)** Distribution of the log(*p*-value) based on 1, 000 independent replicates of the permutation analysis for MDM2 and cAMP, respectively. The cutoff of 8 is indicated by the dashed blue line.

**Table 2.** – Results of CONSTRUCT for each case study.

| Protein | Coordinates | Patch detected | log(p-value) | Optimal length (Å) | Signification of the patch |
| --- | --- | --- | --- | --- | --- |
| Cytochrome c | Cα | Yes | 23.97 | 12 | Binding site with heme molecule |
|  | Side chain | Yes | 15.99 | 13 |  |
| MDM2 | Cα | Yes | 15.82 | 9 | Partially overlaps with p53 interaction surface |
|  | Side chain | Yes | 15.07 | 10 |  |
| KEAP1 (propeller) | Cα | Yes | 65.43 | 19 | Interaction surface with Nrf2 |
|  | Side chain | Yes | 74.63 | 14 |  |
| Myoglobin | Cα | Yes | 48.01 | 17 | Interaction surface with Cu(II) |
|  | Side chain | Yes | 41.86 | 20 |  |
| CFTR | Cα | Yes | 285.39 | 19 | NBD1 domain, including several amino acid sites targeted by drugs |
|  | Side chain | Yes | 249.92 | 20 |  |
| cAMP-dependent protein kinase A | Cα | Yes | 62.17 | 14 | Active kinase core of the protein |
|  | Side chain | Yes | 78.87 | 17 |  |
| DHFR | Cα | Yes | 71.69 | 15 | Binding site with NADPH (and pyrimethamine antimalarial drug) |
|  | Side chain | Yes | 72.82 | 16 |  |
| DHPS | Cα | Yes | 66.32 | 10 | Binding site with sulfa derivatives (such as sulfadoxine) |
|  | Side chain | Yes | 60.62 | 19 |  |
| Mitogen-activated protein kinase 1 | Cα | Yes | 60.67 | 16 | Catalytic site of the protein |
|  | Side chain | Yes | 50.73 | 17 |  |
| SGLT1 | Cα | Yes | 74.46 | 20 | Binding site with glucose |
|  | Side chain | Yes | 88.76 | 20 |  |
| Torsin-1B | Cα | Yes | 122.16 | 20 | Binding site with ATP |
|  | Side chain | Yes | 106.53 | 19 |  |
| YddG | Cα | Yes | 69.60 | 14 | Core of the transporter, binding site with aromatic amino acids |
|  | Side chain | Yes | 54.67 | 16 |  |
| GDP-mannose transporter 1 | Cα | Yes | 66.20 | 17 | Core of the transporter, binding site with GMP |
|  | Side chain | Yes | 54.76 | 8 |  |
| HRas | Cα | Yes | 36.36 | 16 | Biding site with its effectors |
|  | Side chain | Yes | 29.44 | 17 |  |

Permutations of the site-specific substitution rates resulted in an exponential decay of the log(*p*-value). We observed that only 2.2% and 0.31% of the permuted datasets were associated with a log(*p*-value) ≥ 8 for MDM2 and cAMP, respectively, confirming that the threshold of 8 is very conservative, regardless of whether the proteins are small or relatively large (**Figure 2C** and **2F**). This stringent threshold ensures the reliability of detecting true spatial correlations in site-specific substitution rates that indicate functional or structurally important amino acid patches.

### CONSTRUCT identifies functional amino acid patches

We then applied CONSTRUCT to nine other proteins. Here, we focus on two case studies: the CFTR protein, which is responsible for chloride ion flux and whose mutations cause cystic fibrosis (27), and the transmembrane protein YddG, which is found in bacteria and is involved in the export of aromatic amino acids (28). The results for the other case studies are described in the supplementary material (**Supplementary Data S2**).

Rate4Site was first run on a dataset of 351 CFTR orthologous sequences and revealed that the most conserved amino acid sites were uniformly distributed throughout the tertiary structure of CFTR (**Figure 3A**). Of note, these conserved sites included Gly178, Ser549, and Gly551, which are targets of ivacaftor (used to treat cystic fibrosis) (29–31), Gln493, which is thought to be important for ATP hydrolysis (32), and Phe508, which is associated with the major mutation that confers cystic fibrosis (Phe508 deletion) (32). CONSTRUCT was then run on the same dataset. A patch of conserved amino acid sites was detected (log(*p*-value) = 249.92) at an optimal distance of 20 Å (**Table 2**). This patch covered a large part of one of the two NBD domains (NBD1), critical for regulation of channel activity and binding and hydrolysis of ATP (**Figure 3B** and **3C**) (33). The patch included the functional amino acid sites mentioned above, but also Trp496, which is close to Phe508 and reduces the amount of CFTR when mutated to arginine (34), and Leu558, which induces delayed compaction of nascent NBD1 when mutated to serine (35).

**Figure 3.**
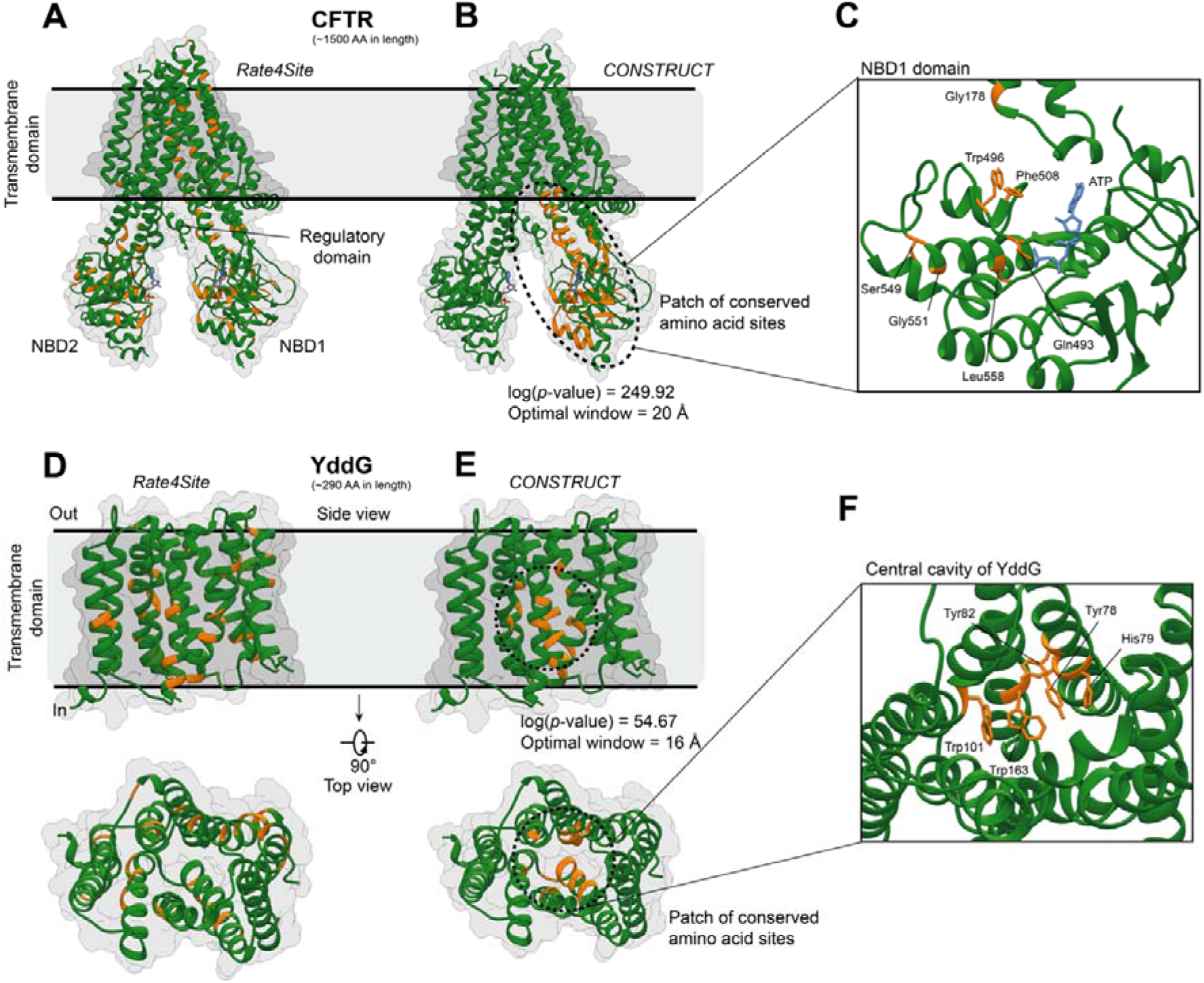
– Identification of patches of functionally validated amino acid sites in CFTR and YddG proteins. Location of the 10% most conserved amino acid sites (colored in orange) in the tertiary structure of CFTR and YddG according to **(A** and **D)** Rate4Site and **(B** and **E)** CONSTRUCT. The functional domains of CFTR and YddG are annotated. **(C** and **F)** Zoom on some amino acid sites in the NBD1 domain of CFTR (which are essential for ATP hydrolysis or cause cystic fibrosis) and in the central channel of YddG (which are reported to be critical for transport activities), which belong to the patches of conserved amino acid sites detected by CONSTRUCT.

**Figure 3.**
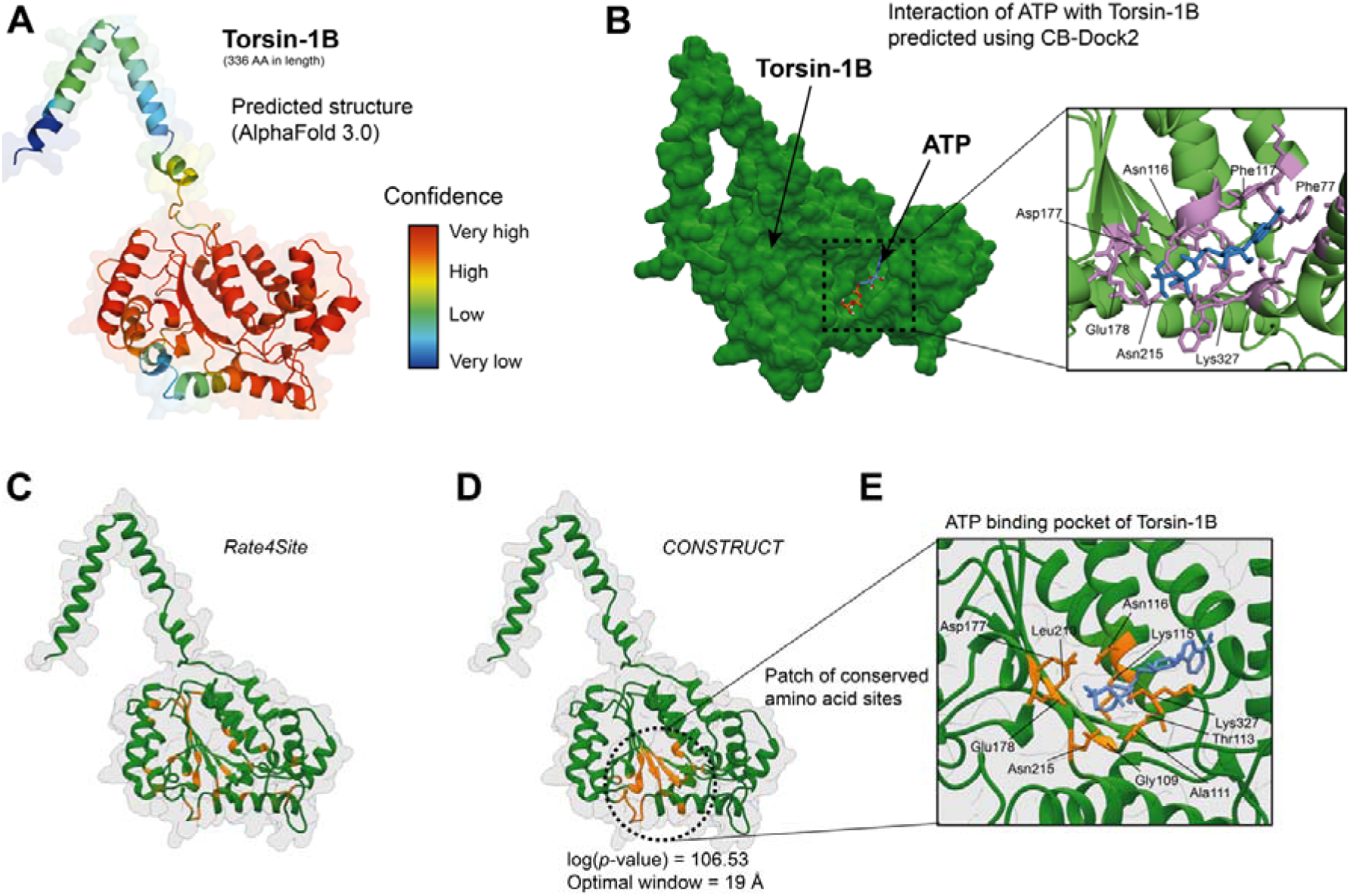
– Identification of patch of conserved amino acid sites using a predicted tertiary structure. **(A)** Predicted tertiary structure of the Torsin-1B protein. The color scale corresponds to the confidence of the prediction for each amino acid site according to AlphaFold 3.0. **(B)** Predicted interaction of ATP with Torsin-1B. Torsin-1B and ATP molecule are shown as surface and stick, respectively. A zoom of the ATP binding pocket shows likely important amino acid sites of Torsin-1B involved in the interaction. **(C** and **D)** Location of the 10% most conserved amino acid sites (colored in orange) in the tertiary structure of Torsin-1B according to Rate4Site and CONSTRUCT, respectively. **(E)** Zoom on some amino acid sites in the predicted ATP-binding pocket of Torsin-1B, which belong to the patch of conserved amino acid sites detected by CONSTRUCT.

We next focused on YddG to test whether CONSTRUCT is able to capture patches of conserved amino acid sites in the context of transporter proteins. Here we used a dataset consisting of 366 YddG orthologous sequences. Again, the most conserved amino acid sites detected by Rate4Site were distributed throughout the protein structure (**Figure 3D**). A few amino acid sites were located in the channel of the transporter, and only Trp101 – reported to be important for transport activity – had its side chain oriented towards the channel (28). CONSTRUCT identified a patch of conserved amino acid sites in the core of the transporter (log(*p*-value) = 54.67) at an optimal distance of 16 Å (**Figure 3E** and **Table 2**). The patch included Trp101, but also Tyr78, His79, Tyr82, and Trp163 (**Figure 3F**), all of which have been reported to be critical for transport activities as evidenced by site-directed mutagenesis (28).

In other case studies, a patch of conserved amino acid sites was systematically detected, in contrast to the broad distribution of conserved amino acid sites proposed by Rate4Site (**Table 2** and **Supplementary Data S2**). All detected patches were associated with experimentally verified functional regions. Overall, the results showed that CONSTRUCT can effectively identify patches of amino acid sites associated with functional constraints.

### An algorithm compatible with predicted structures using AlphaFold

To run CONSTRUCT, the tertiary structure of the protein of interest is required. However, only a limited number of protein structures have been solved experimentally. To address this, artificial intelligence-based approaches, such as Google DeepMind’s AlphaFold, have been developed to accurately predict protein tertiary structures. We wanted to see if CONSTRUCT could precisely identify functional regions based on predicted structures. We studied the protein Torsin-1B, a member of the AAA+ (ATPases Associated with diverse cellular Activities) superfamily, which is involved in several cellular processes, including proper protein folding, maintenance of cellular homeostasis, and possibly the secretory pathway (36). The predicted tertiary structure of Torsin-1B suggested a central domain responsible for ATP binding and hydrolysis (**Figure 4A**), which drives the conformational changes required for its function. It has been reported that Glu178Gln results in a loss of ATPase activity while enhancing the interaction with TOR1AIP2 (36). The interaction with ATP was verified using CB-Dock2 (default parameters) (37), which predicted the interaction to occur within the protein cavity, involving Glu178 and 21 other amino acid sites, with an estimated binding affinity of -8.8 kcal/mol (**Figure 4B**).

Rate4Site was first run on a dataset of 299 Torsin-1B orthologous sequences, revealing that the most conserved amino acid sites were uniformly distributed throughout the tertiary structure of Torsin-1B (**Figure 4C**). CONSTRUCT was then run on the same dataset. A patch of conserved amino acid sites was detected (log(*p*-value) = 106.53) at an optimal distance of 19 Å (**Table 2**). This patch of conserved amino acid sites covered the predicted ATP interaction surface of Torsin-1B (**Figure 4D**). The patch included the amino acid sites Gly109, Ala111, Thr113, Lys115, Asn116, Asp177, Glu178, Leu213, Asn215, and Lys327, all of which are involved in the predicted ATP interaction (**Figure 4B** and **4E**). Notably, the patch contained Glu178 (ranked as the second most conserved site based on spatially correlated site-specific substitution rates), which is essential for the ATPase activity of Torsin-1B (36). Taken together, these results showed that CONSTRUCT can effectively identify functional regions even when using predicted tertiary structures.

### Comparison between C**α** and side chain orientation

Finally, we aimed to compare the delineation of conserved amino acid patches using either the Cα atoms or the center of mass of the amino acids (taking into account the side chain orientation) for a given protein. Here we illustrate the differences obtained for the Kelch-repeat propeller domain of the KEAP1 protein, which is involved in ubiquitination complexes and regulates the cellular level of its substrate Nrf2 (38), and the GDP-mannose transporter 1 membrane protein, which is essential for the transport of GDP-mannose from the cytosol to the lumen of the Golgi apparatus (39). Again, application of the Rate4Site algorithm revealed that conserved amino acid sites are uniformly distributed across the three-dimensional structures of the KEAP1 propeller and GDP-mannose transporter 1 (**Figure 5A** and **5E**), based on 135 and 366 orthologous sequences, respectively.

**Figure 5.**
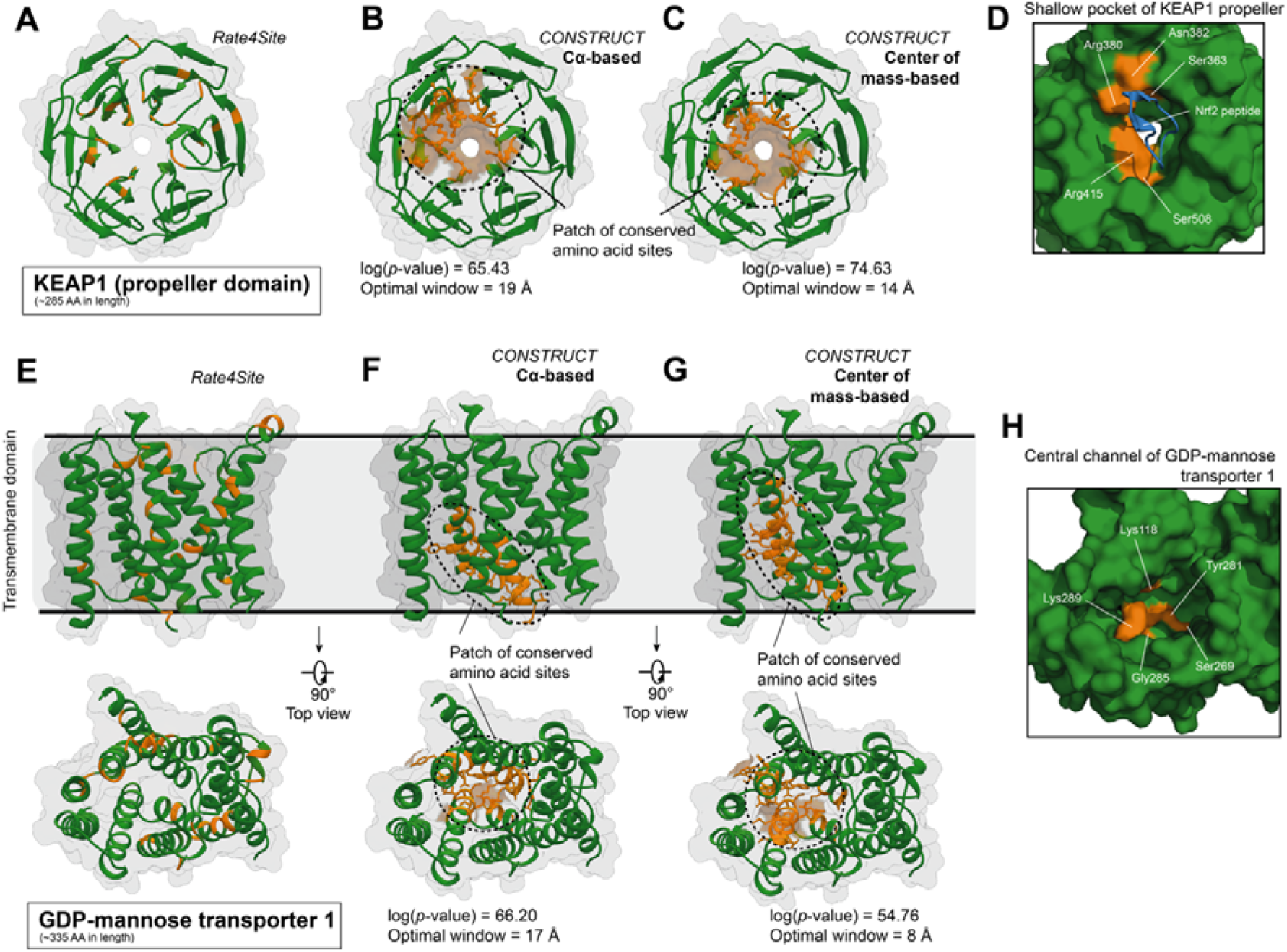
– Comparison of functional patch delineation using either. **C**_α_ **or center of mass of amino acid sites as input coordinates.** Location of the 10% most conserved amino acid sites (colored in orange) in the tertiary structure of KEAP propeller and GDP-mannose transporter 1 according to **(A** and **E)** Rate4Site and CONSTRUCT using either **(B** and **F)** the C_α_ atoms or **(C** and **G)** the center of mass of amino acid sites as input coordinates. **(D** and **H)** Zoom on some amino acid sites in the shallow pocket of KEAP1 propeller (which interact with Nrf2 (shown as cartoon and colored in blue)) and in the central channel of GDP-mannose transporter 1 (which are reported to be critical for transport activities) that belong to the patches of conserved amino acid sites detected by CONSTRUCT.

CONSTRUCT was then applied in parallel using either the Cα atoms or the center of mass of the amino acids. For KEAP1, both strategies identified a conserved patch at the shallow pocket on the surface of the propeller domain, which was experimentally shown to be the interaction surface with Nrf2. Based on the Cα atoms, the patch was delineated with an optimal distance of 19 Å and a log(*p*-value) of 65.43 (**Figure 5B** and **Table 2**). Four amino acids reported to be essential for interaction with Nrf2 were captured: Ser363, Arg380, Asn382, and Arg415 (38). Using the centers of mass of the amino acids, the patch was delineated with an optimal distance of 14 Å and a log(*p*-value) of 74.63 (**Figure 5C** and **Table 2**). Thus, the correlation strength was stronger when based on centers of mass. In addition, the mapping of the most conserved amino acid sites onto the structure revealed a better capture of the shallow pocket compared to the data obtained with the Cα atoms. Furthermore, in addition to the four amino acids mentioned above, we also captured Ser508, which is also reported to be important for the interaction with Nrf2 (38) (**Figure 5D**).

The same strategy was applied to the GDP-mannose transporter 1 protein. Using both the Cα atoms and the centers of mass, a conserved amino acid patch was observed within the transporter channel, indicating the interaction zone with the transporter’s target molecule. With the Cα atoms, the patch was delineated with an optimal distance of 17 Å and a log(*p*-value) of 66.20 (**Figure 5F** and **Table 2**). Two amino acids important for the interaction with GDP-mannose or GMP substrates were located within this patch: Tyr281 and Gly285 (40). In terms of centers of mass, the patch was delineated with an optimal distance of 8 Å and a log(*p*-value) of 54.76 (**Figure 5G** and **Table 2**). Although the correlation strength was lower, the optimal distance was significantly shorter than for the Cα atoms, suggesting that considering side chain orientation allowed for a faster capture of the patch. This patch included the two previously mentioned amino acids as well as Lys118, Lys289, and Ser269, which are also involved in the interaction with GDP-mannose or GMP (40) (**Figure 5H**).

Taken together, these results showed that including the orientation of amino acid side chains in the search for conserved amino acid patches is complementary to the Cα-based analysis and can even improve the delineation of patches in specific structural contexts.

## Discussion

Positive selection and purifying selection are two evolutionary mechanisms that affect the frequency of mutations in populations (1). Positive selection favors beneficial mutations that provide an adaptive advantage to the organism, thereby increasing their frequency over time (1, 3). This process often leads to the emergence of new functions or the improvement of existing ones. In contrast, purifying selection eliminates deleterious mutations that could harm the organism’s survival or reproduction (1, 3). This mechanism helps to maintain essential functions. Consequently, purifying selection is particularly important for identifying functional regions within proteins, because sites under strong purifying selection are usually critical for the function or structure of the protein (3, 14). These sites tend to evolve slowly because mutations that occur there are often deleterious and thus quickly eliminated by natural selection.

Many phylogenetic methods have been developed to identify slowly evolving amino acid sites (5, 11–14). However, the most commonly used methods, such as Rate4Site (14), ignore the spatial correlation of site-specific substitution rates and/or have various limitations. More recently, relatively innovative methods such as GP4Rate and FuncPatch have been developed (20, 21), but they also have their own drawbacks. In addition to being inaccessible, the spatial correlation of site-specific substitution rates is based solely on the polypeptide backbone, which can be a hindrance in structures where the orientation of side chains is critical for protein function, especially in membrane transporters. To address these limitations, we have developed CONSTRUCT, an intuitive software with a graphical interface to make it accessible to biologists and researchers not familiar with bioinformatics. Users only need to provide an alignment of orthologous sequences, a corresponding 3D structure in PDB format, and the structural domains of interest.

CONSTRUCT is based on a simple strategy, which explains the relatively short computation times. Site-specific substitution rates are first calculated using the Rate4Site strategy and then weighted according to their environment. Unlike most other approaches, the distance that maximizes the spatial correlation of site-specific substitution rates is determined directly by calculating all possible rates for an amino acid site at a distance between 1 and 20 Å. Defining the center of mass of the most conserved amino acid sites as the focal point, a simple comparison of the Euclidean distances from this point to the most conserved amino acid sites and to the rest of the “less conserved” amino acid sites confirms the existence of a cluster of conserved amino acid sites. Through a random permutation approach of the site-specific substitution rates, designed to destroy the spatial correlation of the rates, we have defined a very conservative threshold beyond which a patch of conserved amino acid sites is considered to be present, regardless of protein size.

Through a series of 14 case studies, covering a wide range of proteins of different sizes and levels of interspecies conservation, we showed CONSTRUCT’s ability to identify patches of amino acid sites important for protein function and/or structure. We also observed that in all cases, experimentally confirmed functional amino acid sites were consistently captured within the patches. These results confirm that CONSTRUCT can be used to target candidate amino acid sites for directed mutagenesis efforts to elucidate protein function. With the advent of artificial intelligence, we also confirmed that the use of structures generated with AlphaFold (DeepMind, Google) (22) allowed the determination of functional amino acid patches. Finally, we showed that examining the spatial correlation of site-specific substitution rates based on the center of mass of amino acid sites (a proxy for side chain orientation), rather than the peptide backbone, could have a significant impact on the definition of conserved amino acid patches for certain three-dimensional folds. While the overall results were similar between the two approaches, we found that the capture of the conserved amino acid patch within the propeller domain of KEAP1 and within the GDP-mannose transporter 1 membrane transporter was much more precise. While the approaches can be complementary, we believe that considering side chain orientation in the calculation of spatially correlated site-specific substitution rates better reflects the evolutionary dynamics of proteins, as previously proposed (41).

However, the CONSTRUCT tool has some limitations. In particular, it is recommended that the protein be broken down into structural domains if the protein contains multiple domains with possibly different functions. When running CONSTRUCT on such a protein, each structural domain should have its own conserved amino acid patch. Since the program calculates a single focal point (the center of mass of the most conserved amino acid sites), this would be shared by one or more different domains, leading to a biased statistical calculation. To address this, we have included a boundary system that allows the user to specify the coordinates of the different domains, which will then be analyzed independently. Similarly, the tool is not suitable for working on oligomers composed of multiple structural domains. Finally, it is conceivable that functional and/or structurally important regions of the protein may consist of a few to many amino acid sites, depending on the protein family being studied. However, the strategy implemented here is based on a fixed difference between the top 10% most conserved amino acid sites and the rest of the amino acid sites. A possible optimization would be to allow the user to define this threshold or to dynamically calculate this percentage using advanced mathematical models. Additional studies on the proportion of highly conserved amino acid sites in space across protein families should be performed before implementing such functionality.

In conclusion, CONSTRUCT is a simple and intuitive software that allows the determination of regions within a protein that are subject to strong purifying selection, often related to the physiological function of the protein. It can also suggest candidate amino acid sites for functional elucidation via directed mutagenesis approaches.

## References

1. Zhang, J. and Yang, J.-R. (2015) Determinants of the rate of protein sequence evolution. Nat Rev Genet, 16, 409–420.

2. Kimura, M. and Ohta, T. (1974) On some principles governing molecular evolution. Proc Natl Acad Sci U S A, 71, 2848–2852.

3. Echave, J., Spielman, S.J. and Wilke, C.O. (2016) Causes of evolutionary rate variation among protein sites. Nat Rev Genet, 17, 109–121.

4. Yang, Z. (1996) Among-site rate variation and its impact on phylogenetic analyses. Trends Ecol Evol, 11, 367–372.

5. Capra, J.A. and Singh, M. (2007) Predicting functionally important residues from sequence conservation. Bioinformatics, 23, 1875–1882.

6. Franzosa, E.A. and Xia, Y. (2009) Structural determinants of protein evolution are context-sensitive at the residue level. Mol Biol Evol, 26, 2387–2395.

7. Nevin Gerek, Z., Kumar, S. and Banu Ozkan, S. (2013) Structural dynamics flexibility informs function and evolution at a proteome scale. Evol Appl, 6, 423–433.

8. Yeh, S.-W., Liu, J.-W., Yu, S.-H., Shih, C.-H., Hwang, J.-K. and Echave, J. (2014) Site-specific structural constraints on protein sequence evolutionary divergence: local packing density versus solvent exposure. Mol Biol Evol, 31, 135–139.

9. Yang, Z. (1994) Maximum likelihood phylogenetic estimation from DNA sequences with variable rates over sites: approximate methods. J Mol Evol, 39, 306–314.

10. Yang, Z., Nielsen, R., Goldman, N. and Pedersen, A.M. (2000) Codon-substitution models for heterogeneous selection pressure at amino acid sites. Genetics, 155, 431–449.

11. Simon, A.L., Stone, E.A. and Sidow, A. (2002) Inference of functional regions in proteins by quantification of evolutionary constraints. Proc Natl Acad Sci U S A, 99, 2912–2917.

12. Innis, C.A., Anand, A.P. and Sowdhamini, R. (2004) Prediction of functional sites in proteins using conserved functional group analysis. J Mol Biol, 337, 1053–1068.

13. Nimrod, G., Glaser, F., Steinberg, D., Ben-Tal, N. and Pupko, T. (2005) In silico identification of functional regions in proteins. Bioinformatics, 21 **Suppl 1**, i328–337.

14. Pupko, T., Bell, R.E., Mayrose, I., Glaser, F. and Ben-Tal, N. (2002) Rate4Site: an algorithmic tool for the identification of functional regions in proteins by surface mapping of evolutionary determinants within their homologues. Bioinformatics, 18 **Suppl 1**, S71–77.

15. Madabushi, S., Yao, H., Marsh, M., Kristensen, D.M., Philippi, A., Sowa, M.E. and Lichtarge, O. (2002) Structural clusters of evolutionary trace residues are statistically significant and common in proteins. J Mol Biol, 316, 139–154.

16. Panchenko, A.R., Kondrashov, F. and Bryant, S. (2004) Prediction of functional sites by analysis of sequence and structure conservation. Protein Sci, 13, 884–892.

17. Suzuki, Y. (2004) Three-dimensional window analysis for detecting positive selection at structural regions of proteins. Mol Biol Evol, 21, 2352–2359.

18. Berglund, A.-C., Wallner, B., Elofsson, A. and Liberles, D.A. (2005) Tertiary windowing to detect positive diversifying selection. J Mol Evol, 60, 499–504.

19. Watabe, T. and Kishino, H. (2013) Spatial distribution of selection pressure on a protein based on the hierarchical Bayesian model. Mol Biol Evol, 30, 2714–2722.

20. Huang, Y.-F. and Golding, G.B. (2014) Phylogenetic Gaussian Process Model for the Inference of Functionally Important Regions in Protein Tertiary Structures. PLoS Comput Biol, 10, e1003429.

21. Huang, Y.-F. and Golding, G.B. (2015) FuncPatch: a web server for the fast Bayesian inference of conserved functional patches in protein 3D structures. Bioinformatics, 31, 523– 531.

22. Abramson, J., Adler, J., Dunger, J., Evans, R., Green, T., Pritzel, A., Ronneberger, O., Willmore, L., Ballard, A.J., Bambrick, J., et al. (2024) Accurate structure prediction of biomolecular interactions with AlphaFold 3. Nature, 630, 493–500.

23. Kussie, P.H., Gorina, S., Marechal, V., Elenbaas, B., Moreau, J., Levine, A.J. and Pavletich, N.P. (1996) Structure of the MDM2 Oncoprotein Bound to the p53 Tumor Suppressor Transactivation Domain. Science, 274, 948–953.

24. Turnham, R.E. and Scott, J.D. (2016) Protein kinase A catalytic subunit isoform PRKACA; History, function and physiology. Gene, 577, 101–108.

25. Moll, U.M. and Petrenko, O. (2003) The MDM2-p53 Interaction. Molecular Cancer Research, 1, 1001–1008.

26. Yang, J., Ten Eyck, L.F., Xuong, N.-H. and Taylor, S.S. (2004) Crystal Structure of a cAMP-dependent Protein Kinase Mutant at 1.26 Å: New Insights into the Catalytic Mechanism. Journal of Molecular Biology, 336, 473–487.

27. Devidas, S. and Guggino, W.B. (1997) CFTR: domains, structure, and function. J Bioenerg Biomembr, 29, 443–451.

28. Tsuchiya, H., Doki, S., Takemoto, M., Ikuta, T., Higuchi, T., Fukui, K., Usuda, Y., Tabuchi, E., Nagatoishi, S., Tsumoto, K., et al. (2016) Structural basis for amino acid export by DMT superfamily transporter YddG. Nature, 534, 417–420.

29. Zhang, Z., Liu, F. and Chen, J. (2018) Molecular structure of the ATP-bound, phosphorylated human CFTR. Proc Natl Acad Sci U S A, 115, 12757–12762.

30. Yarlagadda, S., Zhang, W., Penmatsa, H., Ren, A., Arora, K., Naren, A.P., Khan, F.A.I., Donnellan, C.A., Srinivasan, S., Stokes, D.C., et al. (2012) A Young Hispanic with c.1646G>A Mutation Exhibits Severe Cystic Fibrosis Lung Disease: Is Ivacaftor an Option for Therapy? Am J Respir Crit Care Med, 186, 694–696.

31. Levring, J., Terry, D.S., Kilic, Z., Fitzgerald, G., Blanchard, S.C. and Chen, J. (2023) CFTR function, pathology and pharmacology at single-molecule resolution. Nature, 616, 606–614.

32. Lewis, H.A., Zhao, X., Wang, C., Sauder, J.M., Rooney, I., Noland, B.W., Lorimer, D., Kearins, M.C., Conners, K., Condon, B., et al. (2005) Impact of the ΔF508 Mutation in First Nucleotide-binding Domain of Human Cystic Fibrosis Transmembrane Conductance Regulator on Domain Folding and Structure*. Journal of Biological Chemistry, 280, 1346– 1353.

33. Duffieux, F., Annereau, J.P., Boucher, J., Miclet, E., Pamlard, O., Schneider, M., Stoven, V. and Lallemand, J.Y. (2000) Nucleotide-binding domain 1 of cystic fibrosis transmembrane conductance regulator production of a suitable protein for structural studies. Eur J Biochem, 267, 5306–5312.

34. Thibodeau, P.H., Brautigam, C.A., Machius, M. and Thomas, P.J. (2005) Side chain and backbone contributions of Phe508 to CFTR folding. Nat Struct Mol Biol, 12, 10–16.

35. Shishido, H., Yoon, J.S., Yang, Z. and Skach, W.R. (2020) CFTR trafficking mutations disrupt cotranslational protein folding by targeting biosynthetic intermediates. Nat Commun, 11, 4258.

36. Rose, A.E., Zhao, C., Turner, E.M., Steyer, A.M. and Schlieker, C. (2014) Arresting a Torsin ATPase reshapes the endoplasmic reticulum. J Biol Chem, 289, 552–564.

37. Liu, Y., Yang, X., Gan, J., Chen, S., Xiao, Z.-X. and Cao, Y. (2022) CB-Dock2: improved protein-ligand blind docking by integrating cavity detection, docking and homologous template fitting. Nucleic Acids Res, 50, W159–W164.

38. Canning, P., Sorrell, F.J. and Bullock, A.N. (2015) Structural basis of Keap1 interactions with Nrf2. Free Radic Biol Med, 88, 101–107.

39. Abe, M., Hashimoto, H. and Yoda, K. (1999) Molecular characterization of Vig4/Vrg4 GDP-mannose transporter of the yeast Saccharomyces cerevisiae. FEBS Lett, 458, 309–312.

40. Parker, J.L. and Newstead, S. (2017) Structural basis of nucleotide sugar transport across the Golgi membrane. Nature, 551, 521–524.

41. Marcos, M.L. and Echave, J. (2015) Too packed to change: side-chain packing and site-specific substitution rates in protein evolution. PeerJ, 3, e911.

