## Supplementary Data for "CONSTRUCT: an algorithmic tool for identifying functional or structurally important regions in protein tertiary structure"

**Supplementary Data S1 – Description of the proteins investigated in this study.**

**Cytochrome c (Uniprot: P99999, PDB ID: 1J3S)**

Cytochrome c is an essential component of the mitochondrial electron transport chain, a key pathway in cellular respiration. Its primary function is to transfer electrons between complex III (cytochrome *bc1* complex) and complex IV (cytochrome c oxidase), which is critical for the generation of ATP through oxidative phosphorylation (1). In addition to its role in energy production, cytochrome c is also involved in intrinsic apoptosis pathways. During apoptosis, cytochrome c is released from the mitochondria into the cytosol where it interacts with Apaf-1 (apoptotic protease activating factor 1) and caspase-9 to initiate apoptosome formation and activate the apoptotic cascade that executes the cell death program (2, 3). Cytochrome c is located in the intermembrane space of the mitochondria. This localization allows it to shuttle between complex III and complex IV during electron transfer (4). This protein non-covalently binds a heme group, with the heme iron being the site of electron transfer (5). Key functional amino acids include histidine and methionine, specifically His18 (which coordinates the heme iron and is critical for the electron transfer function) and Met80 (which provides a ligand for the heme iron and stabilizes the heme group) (6). In addition, a series of lysine residues (at amino acid positions 8, 13, 27, 72, 79, and 87) surrounding the heme interaction site form electrostatic interactions with the membrane and with components of the cytochrome *bc1* complex and cytochrome c oxidase (7, 8). These residues contribute to the formation of electron transfer complexes and the stabilization of protein-protein interactions (8). Structurally, cytochrome c is a small globular protein (PDB ID: 1J3S). Its three-dimensional structure is highly conserved across eukaryotic species, with a compact fold that stabilizes the heme group and ensures efficient electron transfer. This compact structure includes a combination of alpha helices and loops that provide a stable environment for the heme group, facilitating its role in electron transport.

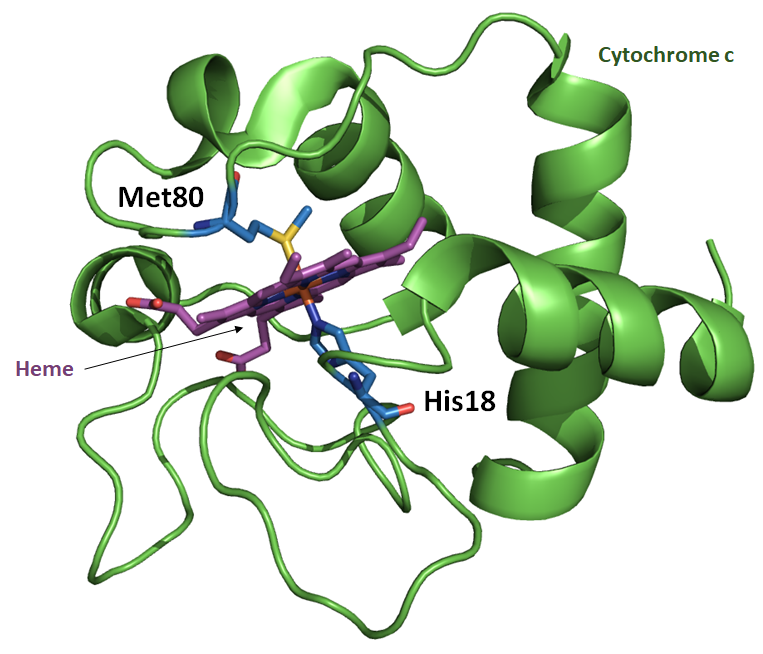

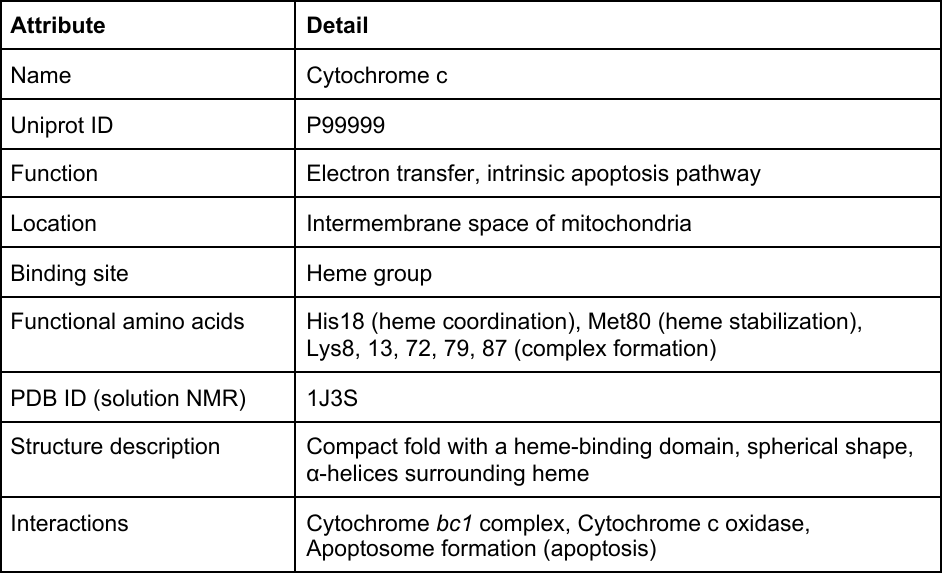

**MDM2 (Uniprot: Q00987, PDB ID: 1YCR)**

The E3 ubiquitin protein ligase MDM2 (mouse double minute 2 homolog) is a key regulator of the p53 tumor suppressor protein, which plays a key role in controlling cell cycle progression and apoptosis (9, 10). The protein targets p53 for proteasomal degradation through ubiquitination, thereby modulating p53 levels and activity within the cell. This regulation is essential for maintaining cellular homeostasis, preventing uncontrolled cell proliferation, and ensuring timely cell death in response to DNA damage or stress. MDM2 is primarily located in the nucleus, but can also shuttle between the nucleus and the cytoplasm (11). This shuttling is essential for its function, as it must interact with p53 in the nucleus and transport it to the cytoplasm for proteasomal degradation (10). The protein contains several functional domains, each of which contributes to its function in ubiquitination and interaction with p53 (12): *i*) an N-terminal p53-binding domain that directly interacts with the transactivation domain of p53, inhibiting its activity and targeting it for degradation; *ii*) a central acidic domain that is involved in interaction with other proteins and can influence the conformation and activity of MDM2; and *iii*) a RING finger domain that is essential for its ubiquitin ligase activity and facilitates the transfer of ubiquitin from an E2 ubiquitin-conjugating enzyme to p53. Critical amino acids include Cys464 within the RING finger domain, which is essential for coordinating zinc ions that stabilize the RING finger structure necessary for ubiquitin ligase activity (13). In addition, specific amino acids in the N-terminal domain (such as Gly58, Glu68, Val75 and Cys77) form the binding site for p53, providing the high-affinity interaction required for effective ubiquitination (14). The three-dimensional structure of MDM2 reveals a complex architecture in which the p53-binding domain forms a deep hydrophobic cleft to accommodate the transactivation domain of p53 (PDB ID: 1YCR). This structure allows MDM2 to effectively regulate p53 levels and activity, thereby influencing cell fate decisions (9). Notably, MDM2 can also interact with the tumor suppressor ARF (alternate reading frame), which can inhibit its activity and lead to p53 stabilization and activation (15).

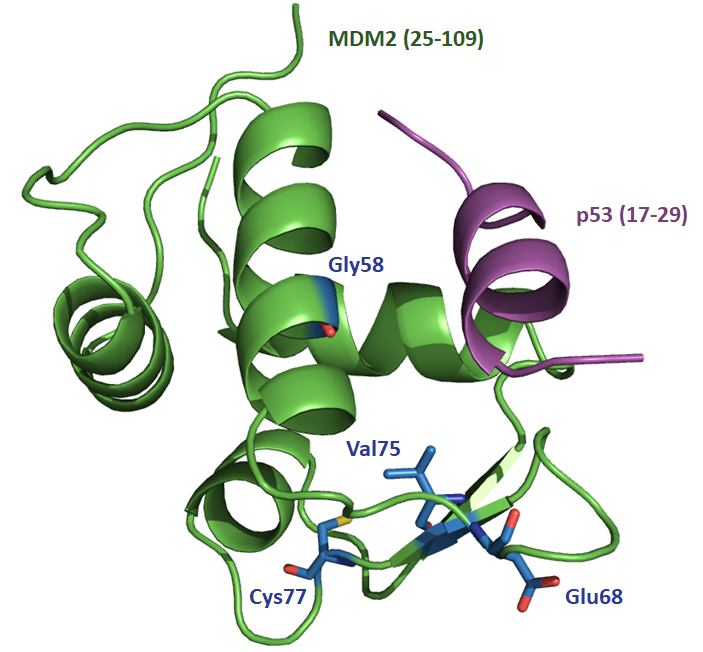

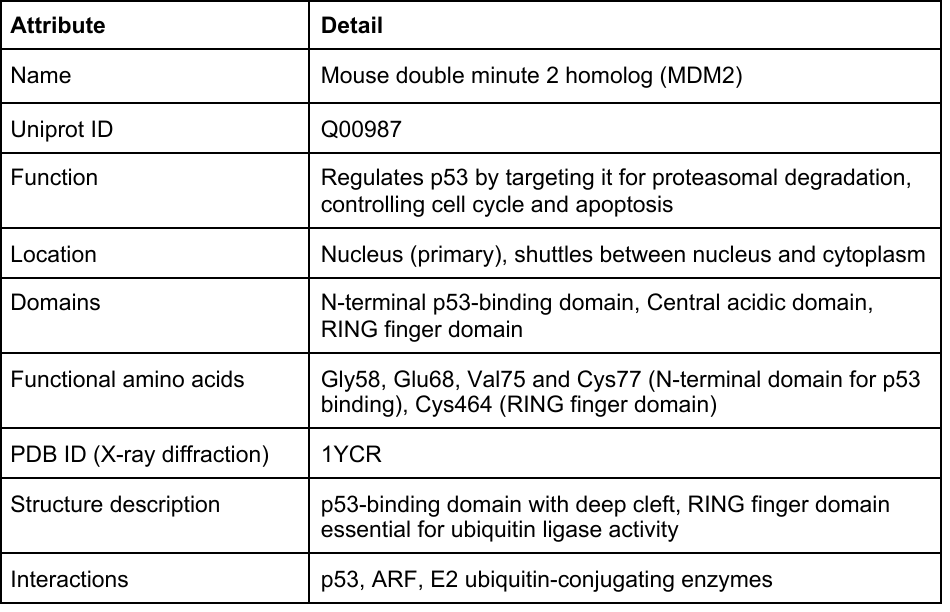

**KEAP1 (Uniprot: Q14145, PDB ID: 2FLU)**

KEAP1 is a key regulator of the nuclear factor erythroid 2-related factor 2 (Nrf2) pathway, which plays an important role in the cellular response to oxidative stress (16). KEAP1 functions as an adaptor protein for the cullin 3 (Cul3) ubiquitin ligase complex, targeting Nrf2 for ubiquitination and subsequent proteasomal degradation under non-stressed conditions (17, 18). This regulation ensures that Nrf2 levels remain low under normal conditions. Upon exposure to oxidative stress, KEAP1 undergoes conformational changes that prevent it from targeting Nrf2 for degradation, allowing Nrf2 to translocate to the nucleus and activate the expression of antioxidant response element (ARE)-dependent genes (16, 19). KEAP1 is located in the cytoplasm where it forms complexes with Nrf2 and Cul3. It can also shuttle between the cytoplasm and the nucleus, although its primary function is in the cytoplasm. KEAP1 consists of: *i*) a Broad-Complex, Tramtrack and Bric-a-brac (BTB) domain that is essential for homodimerization and interaction with Cul3; *ii*) an IVR (Intervening Region) domain that connects the BTB and Kelch domains and plays a role in sensing oxidative stress; and *iii*) a Kelch domain containing six kelch-repeats that form a β-propeller structure (20, 21). This domain is responsible for binding to Nrf2. Several essential amino acid sites have been reported in KEAP1, including Cys151, Cys273, and Cys288, which are critical for sensing oxidative and electrophilic stress (16), and Ser363, Arg380, Asn382, Arg415, Arg483 and Ser508, which provide high-affinity interaction with Nrf2 for effective ubiquitination (22).

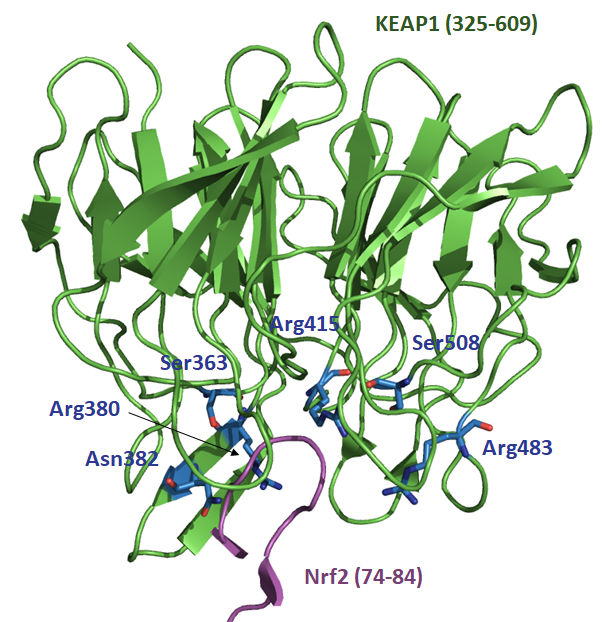

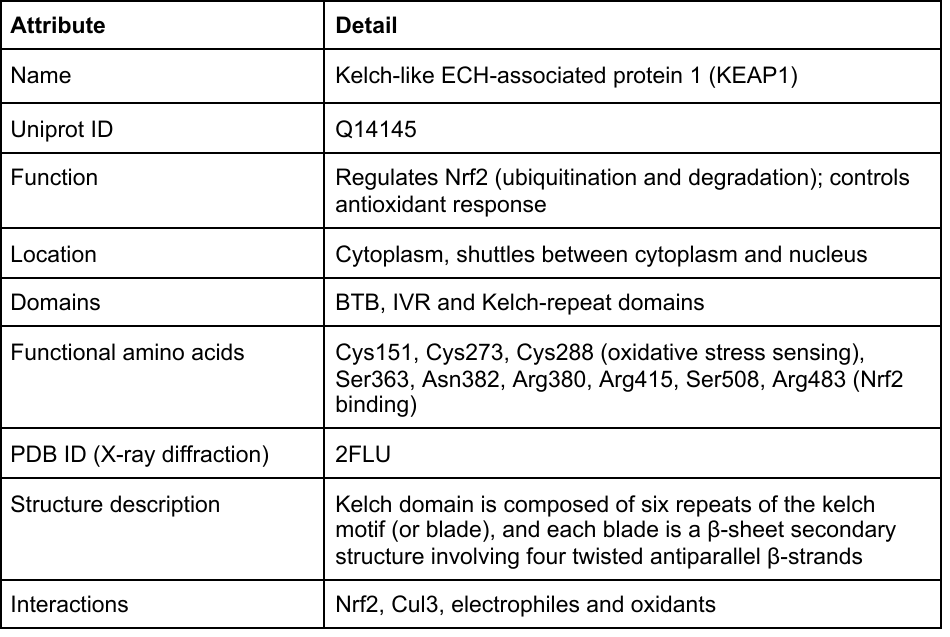

**DHFR (Uniprot: A7UD81, PDB ID: 3QGT)**

*Plasmodium falciparum* dihydrofolate reductase (DHFR) is a critical enzyme in the folate pathway, which is essential for DNA synthesis, repair and methylation. DHFR catalyzes the reduction of dihydrofolate (DHF) to tetrahydrofolate (THF) using NADPH as a cofactor. THF acts as a carrier of single carbon units in various metabolic reactions, including the synthesis of purines, thymidylate, and certain amino acids. Inhibition of DHFR leads to a depletion of THF and subsequently hinders DNA synthesis, making DHFR a primary target for antimalarial drugs such as pyrimethamine (23). DHFR is located in the cytoplasm of *P. falciparum* cells. The enzyme is part of the bifunctional enzyme DHFR-TS (thymidylate synthase) complex, which contains both dihydrofolate reductase and thymidylate synthase activities in a single polypeptide chain. The DHFR enzyme consists of an N-terminal domain containing the DHFR activity and a C-terminal domain containing the thymidylate synthase (TS) activity (24). The enzyme acts as a homodimer, with each monomer contributing to the overall catalytic activity of the enzyme complex (25). Key functional amino acids include *i*) Asp54, which is critical for binding the substrate DHF (26); *ii*) Asn51, which plays a role in substrate and cofactor binding (27); and *iii*) Phe58, which is involved in maintaining the structural integrity of the enzyme (23). The three-dimensional structure of *P. falciparum* DHFR (PDB ID: 3QGT) shows a highly conserved fold typical of DHFR enzymes, with a central eight-stranded β-sheet flanked by α-helices (28). The active site is located at the interface of the β-sheet and one of the α-helices, where it accommodates the substrate (DHF) and the cofactor (NADPH). Mutations in the *dhfr* gene can confer resistance to antifolate drugs such as pyrimethamine and cycloguanil (26–28). These mutations alter the binding affinity of the enzyme for the inhibitors without significantly affecting the binding of the natural substrate or cofactor.

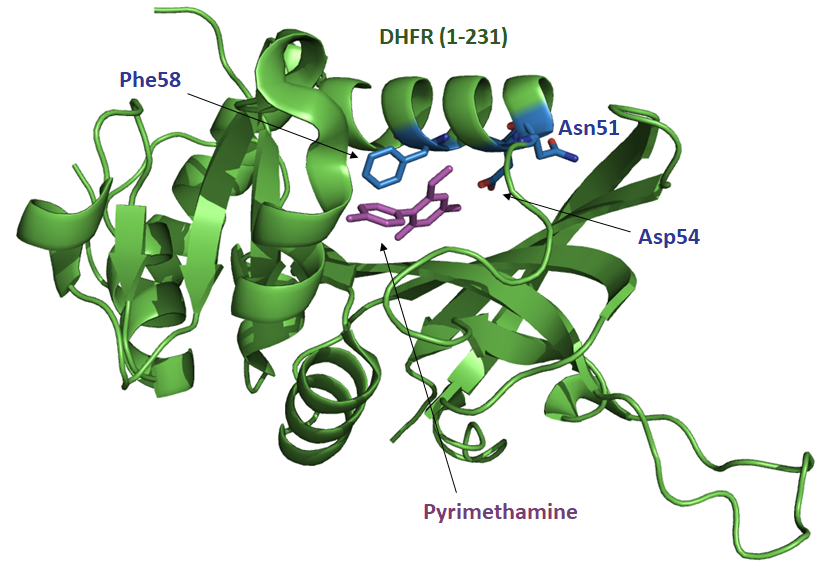

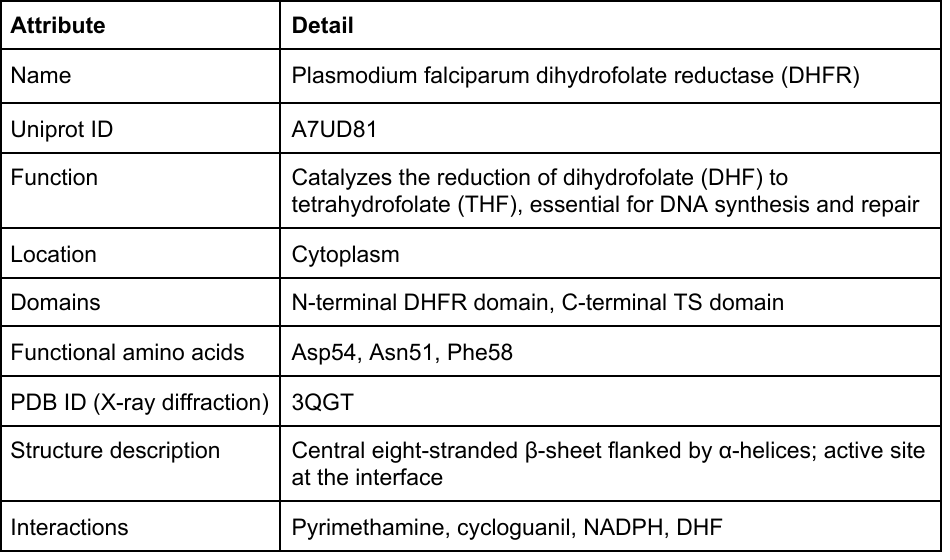

**Myoglobin (Uniprot: P02144, PDB ID: 3RGK)**

Myoglobin is a small heme-containing protein that is primarily responsible for storing and transporting oxygen in muscle tissue (29, 30). It facilitates oxygen diffusion and provides a reserve supply of oxygen, which is critical during periods of high muscular activity. Myoglobin has a high affinity for oxygen, allowing it to effectively capture and release oxygen molecules (31). This function is essential for maintaining the metabolic demands of muscle cells, especially under anaerobic conditions. Myoglobin is located in the cytoplasm of muscle cells, particularly in skeletal and cardiac muscle tissues. Its presence in these tissues allows it to efficiently bind and release oxygen where it is most needed (29, 31). Myoglobin is a globular protein consisting of a single polypeptide chain of 154 amino acids. The protein adopts a globin fold characterized by eight alpha-helices that create a hydrophobic pocket in which the heme group (consisting of an iron (Fe) atom held within a porphyrin ring) is located (32). The protein has been extensively characterized and some functional amino acids have been reported, including Gln91, Ser92, Ala94 and Thr95, which can bind to binuclear Cu(II) for hydrolytic cleavage of the protein (33); and His93, which coordinates directly to the iron atom of the heme group and plays a critical role in structural stability (34). The three-dimensional structure of myoglobin (PDB ID: 3RGK) is highly conserved, with its tightly packed alpha-helices providing a stable environment for the heme group (35). The compact, globular architecture of myoglobin ensures efficient oxygen binding and release, which is critical for its function in muscle tissue. Myoglobin's high affinity for oxygen is a result of the precise positioning of the heme group within the protein.

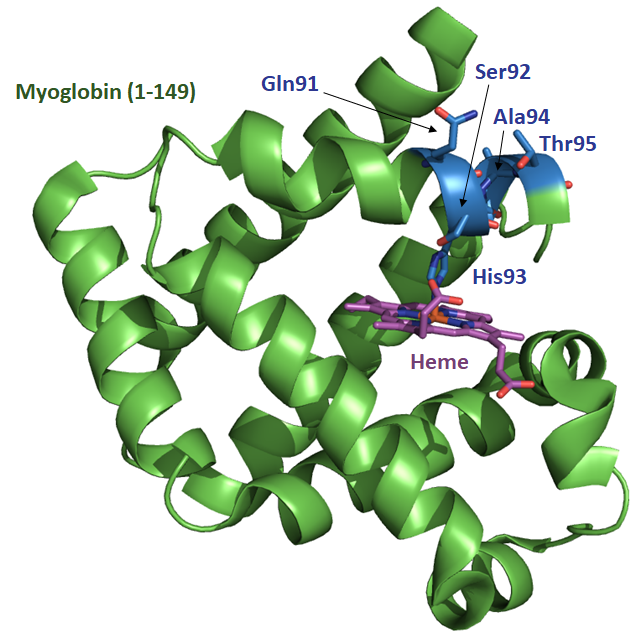

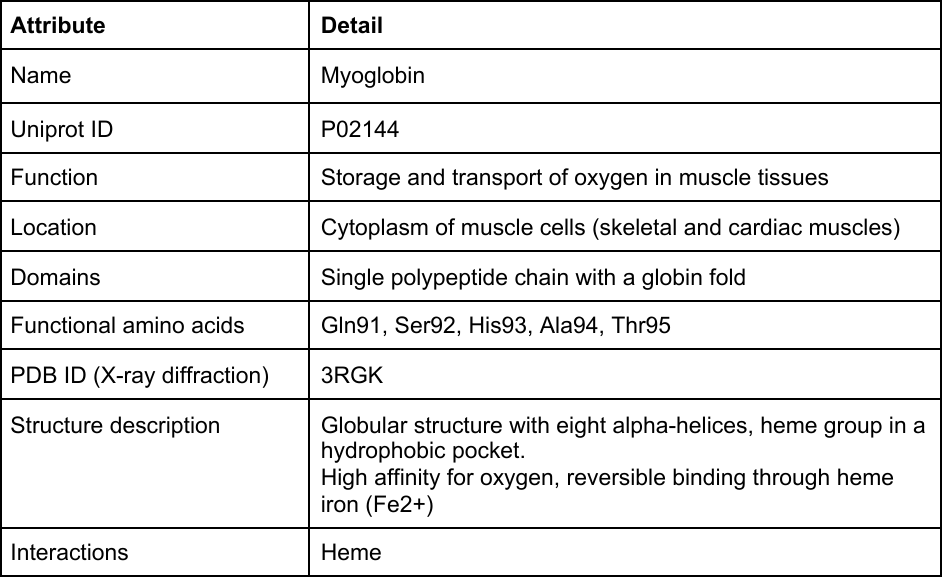

**cAMP-dependent protein kinase A (Uniprot: P17612, PDB ID: 4WB5)**

The cAMP-dependent protein kinase catalytic subunit alpha (PKA Cα) is a key player in the regulation of various cellular processes, including metabolism, gene expression, cell cycle progression, and memory formation (36). PKA Cα is one of the catalytic subunits of the protein kinase A (PKA) holoenzyme, which is activated by cyclic AMP (cAMP). Upon binding cAMP, the regulatory subunits of PKA release the catalytic subunits, allowing them to phosphorylate specific serine and threonine residues on target proteins, thereby modulating their activity (36). PKA Cα is primarily located in the cytoplasm but can translocate to the nucleus upon activation (37). This translocation allows PKA to phosphorylate nuclear substrates, thereby affecting gene expression and other nuclear functions. The protein consists of *i*) an N-terminal domain containing a glycine-rich loop involved in ATP binding; *ii*) a catalytic core containing the small and large lobes of the kinase domain (the small lobe primarily binds ATP, while the large lobe binds the protein substrate); *iii*) an activation segment located within the catalytic core that undergoes conformational changes upon activation; and *iv*) a C-terminal tail containing a hydrophobic motif that interacts with the catalytic core to stabilize the active conformation (36, 38). Several amino acid sites have been reported to be functionally important, such as Lys72, which is involved in ATP binding and stabilization of the phosphate groups (39, 40); Asp166, which is critical for phosphotransferase activity; Thr197, which is located in the activation segment and must be phosphorylated for full catalytic activity (41); and Phe327 and Met231, which are involved in substrate recognition and binding (42, 43). The three-dimensional structure of PKA Cα (PDB ID: 4WB5) has been well characterized, revealing a bilobal kinase fold typical of protein kinases (38). The ATP-binding site is located in a deep cleft between the small and large lobes, with the activation segment playing a critical role in substrate recognition and catalysis (38). PKA is activated by the binding of cAMP to its regulatory subunits, which induces a conformational change leading to the release of the catalytic subunits. Once released, the catalytic subunits phosphorylate target proteins on serine and threonine residues, thereby modulating their function (36). This process is tightly regulated to ensure precise control of cellular signaling pathways.

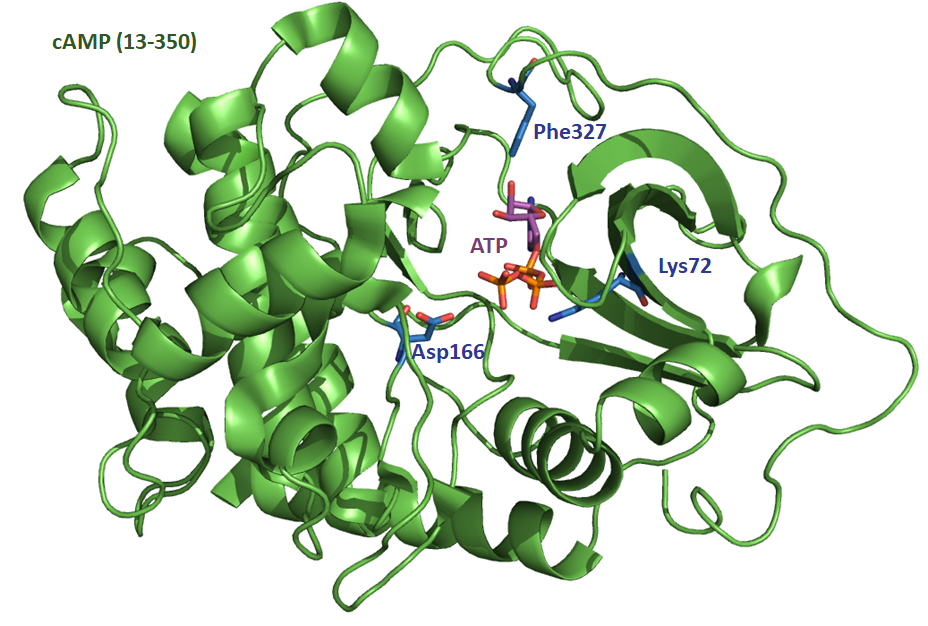

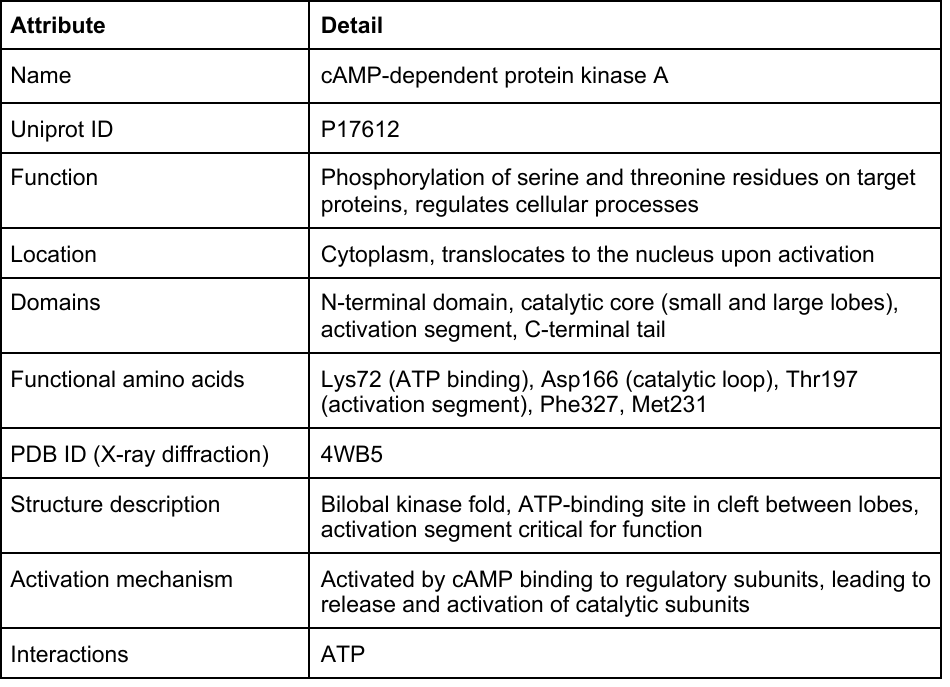

**DHPS (Uniprot: Q25704, PDB ID: 6JWV)**

Dihydropteroate synthase (DHPS) is an enzyme crucial for the folate biosynthesis pathway in *Plasmodium* species, which are the parasites responsible for malaria. DHPS catalyzes the condensation of para-aminobenzoic acid (pABA) with 6-hydroxymethyl-7,8-dihydropterin pyrophosphate to form dihydropteroate (44, 45). This reaction is a key step in the synthesis of dihydrofolate, which is eventually converted into tetrahydrofolate, an essential cofactor for the synthesis of nucleotides and amino acids (44, 45). Inhibition of DHPS disrupts folate metabolism, impairing DNA synthesis and cell division, making it a target for antimalarial drugs (45). DHPS is localized in the cytoplasm of the parasite. It functions within the folate biosynthesis pathway, which operates in the cytoplasmic environment where substrates and cofactors are available. DHPS is constituted of a catalytic domain where the substrate binding and the catalytic reaction occur (46). This domain also includes binding sites for pABA and dihydropterin pyrophosphate) (47). Key functional amino acid sites are Ala437, which is involved in binding dihydropterin pyrophosphate; and Lys540, which plays a critical role in the catalytic mechanism by stabilizing the transition state of the reaction (48, 49). Certain residues (such as Ala581 and Ala613) are also critical for binding sulfa drugs (such as sulfadoxine), which are competitive inhibitors of DHPS (50). Mutations at these sites can confer resistance to these drugs (49, 50). The three-dimensional structure of DHPS from *P. falciparum* has been resolved (PDB ID: 6JWV), providing insights into its function and interaction with inhibitors (46). The enzyme adopts a fold typical of dihydropteroate synthases, with a central beta-sheet flanked by alpha-helices.

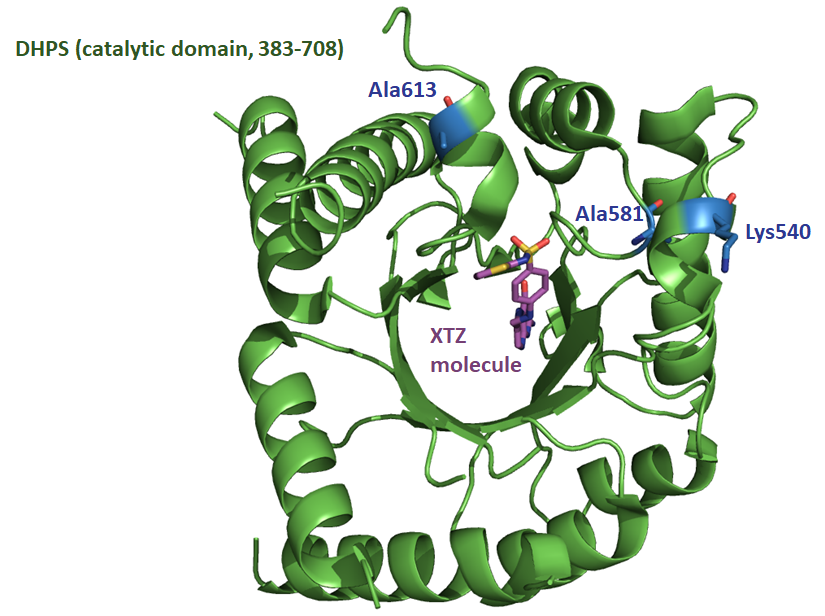

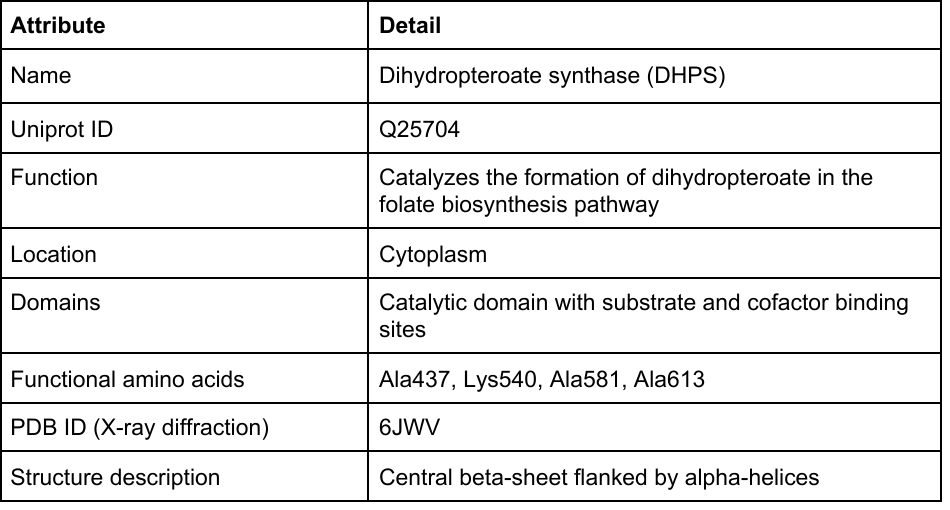

**CFTR (Uniprot: P13569, PDB ID: 8FZQ)**

The cystic fibrosis transmembrane conductance regulator (CFTR) is a vertebrate membrane protein and chloride channel encoded by the *cftr* gene. Its primary function is to transport chloride ions across epithelial cell membranes, which is essential for the regulation of salt and water transport in various tissues (51–53). This regulation is critical for the production of sweat, digestive fluids, and mucus. Dysfunction of CFTR due to genetic mutations leads to cystic fibrosis (CF), a severe genetic disorder that affects the lungs, pancreas and other organs (54–56). CF is characterized by thick, sticky mucus that can obstruct airways and glands, leading to respiratory and digestive complications. CFTR is located in the apical membrane of epithelial cells, particularly in tissues such as the lung, pancreas, intestine, and sweat glands. This apical localization allows CFTR to regulate ion and fluid transport directly at the interface with the external environment or luminal surfaces. CFTR is a large and complex protein consisting of 1,480 amino acids and contains several domains, including *i*) two transmembrane domains (TMDs), which form the channel pore through which chloride ions pass (each TMD consists of six membrane-spanning alpha-helices); *ii*) two nucleotide-binding domains (NBDs), which are critical for the regulation of channel activity and bind and hydrolyze ATP (which controls the opening and closing of the chloride channel); and *iii*) a regulatory (R) domain, which regulates channel activity in response to cellular signals, particularly phosphorylation by protein kinase A (PKA) (51, 57). Key functional amino acids are found in the NBDs and include those involved in ATP binding and hydrolysis. These include the Walker A motif (glycine-rich sequence) (58), the Walker B motif (DEXX box) (59), and residues at positions 458 to 465 that regulate the opening and closing of the chloride channel (60). The three-dimensional structure of CFTR (PDB ID: 1XMJ) shows a complex arrangement with the TMDs forming the chloride channel pore, the cytoplasmic NBDs interacting with ATP, and the R domain modulating channel activity through phosphorylation (61). This structure is critical for the function of CFTR in ion transport and fluid balance in epithelial tissues.

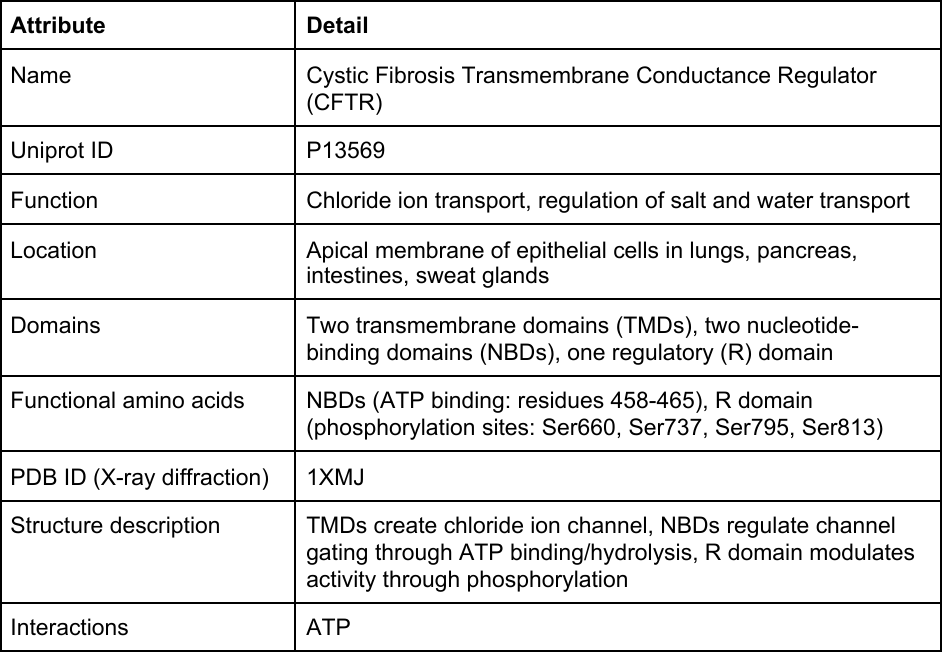

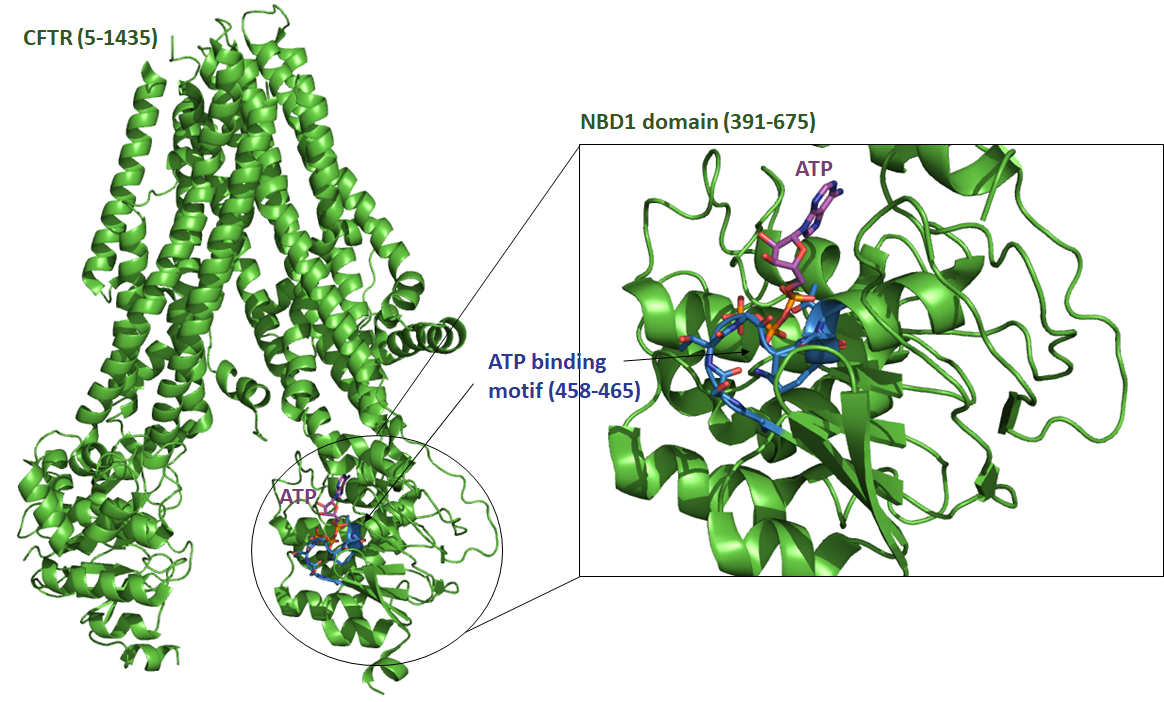

**MAPK1 (Uniprot: P63086, PDB ID: 5UMO)**

Mitogen-activated protein kinase 1 (MAPK1), also known as extracellular signal-regulated kinase 2 (ERK2), is a critical component of the MAP kinase signaling pathway (62, 63). This pathway transduces extracellular signals into the cell and influences various cellular processes such as proliferation, differentiation, development, learning and memory. MAPK1 is activated through phosphorylation by MAP kinase kinase (MEK1/2) in response to growth factors, hormones, and other extracellular stimuli (64). Upon activation, MAPK1 translocates to the nucleus where it phosphorylates various transcription factors and other nuclear substrates to regulate gene expression. MAPK1 is ubiquitously expressed and can be found in both the cytoplasm and the nucleus (65). In its inactive state, it is primarily cytoplasmic, but upon activation, it translocates to the nucleus to exert its effects on target genes (65). MAPK1 consists of: *i*) an N-terminal domain for ATP binding, which is critical for the kinase activity; *ii*) a kinase domain, which contains a highly conserved activation loop that includes residues to be phosphorylated for full activation; *iii*) a C-terminal domain, which is involved in substrate recognition and binding; and *iv*) a nuclear localization signal (NLS), which facilitates the translocation of activated MAPK1 to the nucleus (66). Some amino acid sites were reported to be critical for MAPK1 function, such as Asp147, which is part of the catalytic loop and essential for ATP binding and catalysis (67, 68); or Ala187, Arg189 and Arg192, which participate in substrate recognition (67, 68). The three-dimensional structure of MAPK1 reveals a typical protein kinase fold, comprising a small N-terminal lobe and a larger C-terminal lobe. The ATP-binding site is located in the cleft between these lobes (PDB ID: 5UMO) (66).

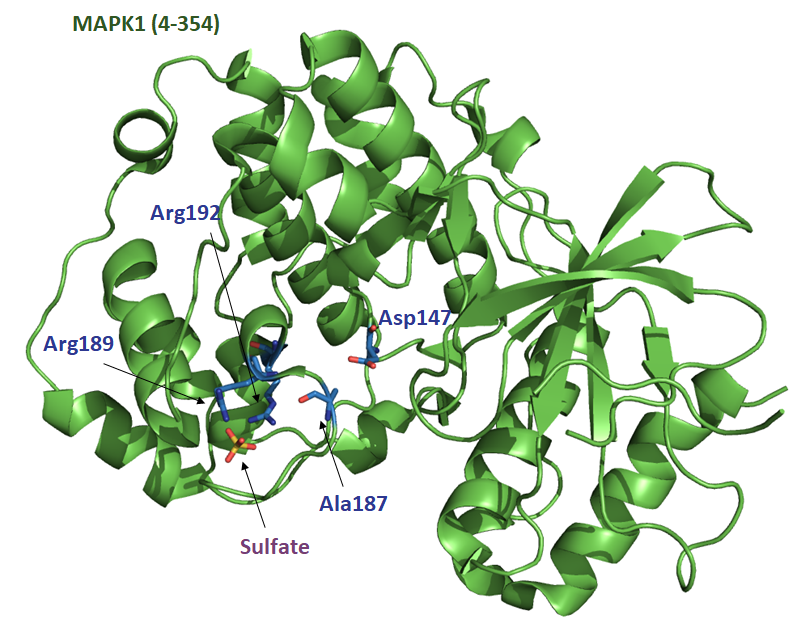

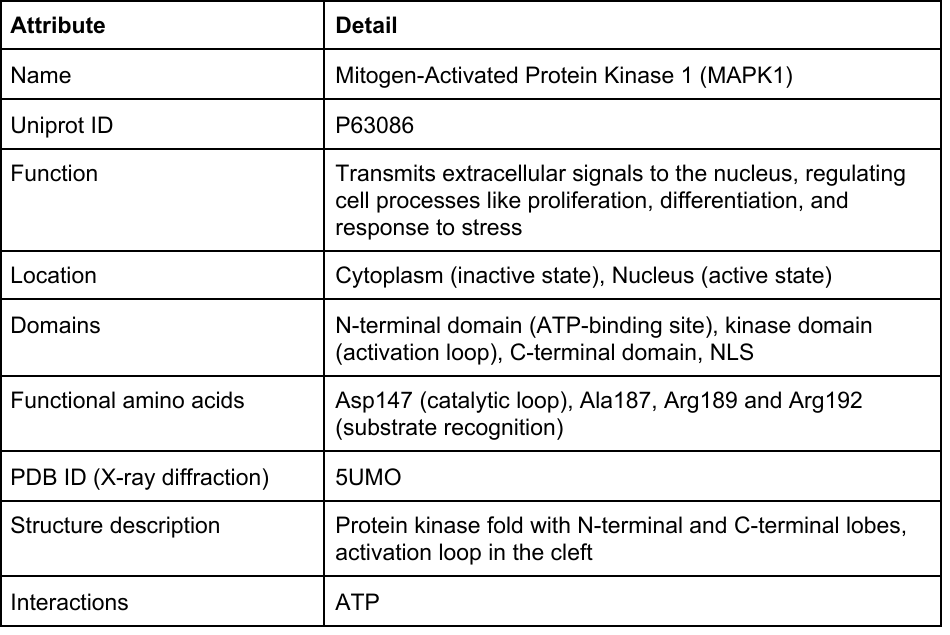

**SGLT1 (Uniprot: P13866, PDB ID: 7SL8)**

Sodium-glucose cotransporter 1 (SGLT1) is a membrane protein that plays a critical role in the active transport of glucose and galactose across the intestinal epithelium (69, 70). SGLT1 uses the electrochemical gradient of sodium ions to drive the uptake of glucose and galactose against their concentration gradients. This process is essential for the absorption of dietary sugars in the small intestine and for renal glucose reabsorption in the proximal tubules of the kidney (69). SGLT1 is primarily located in the apical membrane of enterocytes in the small intestine and in the epithelial cells of the proximal tubules of the kidney (70, 71). This positioning allows it to effectively absorb glucose and galactose from the intestinal lumen and to recover glucose from the glomerular filtrate in the kidney. SGLT1 is a 664-amino acid multi-pass transmembrane protein. SGLT1 has 14 transmembrane helices that form a channel for glucose and sodium transport (72). The loops connecting the transmembrane helices play a role in the protein's regulatory functions and interactions with other cellular components. The N- and C-terminal regions of the protein are involved in regulating the activity of the transporter and its insertion into the membrane (72). Some key amino acid sites have been reported, such as Gly86, Leu452 and Phe453, which are important for glucose binding and recognition (72); or Trp291, which is essential for the uptake of the glucose congener α-methyl-d-glucopyranoside (αMDG) (72). The three-dimensional structure reveals a channel formed by transmembrane helices to which sodium ions and glucose molecules bind and are transported together across the membrane (PDB ID: 7SL8) (72). SGLT1 operates by a symport mechanism in which the simultaneous binding of sodium ions and glucose induces conformational changes that transport both molecules across the membrane. The energy for this process comes from the sodium gradient maintained by the Na+/K+ ATPase pump.

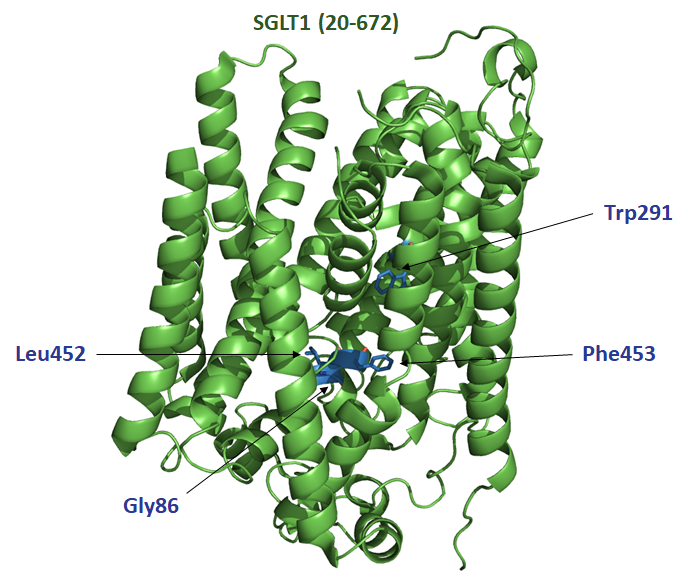

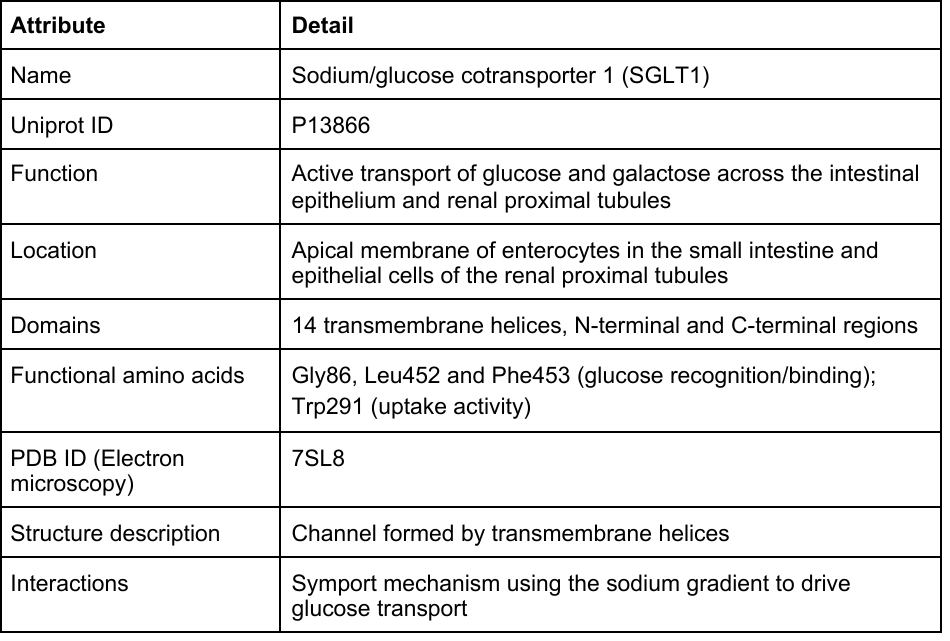

**Torsin-1B (Uniprot: O14657, no PDB ID)**

Torsin-1B is a member of the AAA+ (ATPases Associated with diverse cellular Activities) superfamily, which is involved in several cellular processes, including proper protein folding, maintenance of cellular homeostasis, and possibly the secretory pathway (73, 74). Although its precise functions are not fully understood, Torsin-1B is thought to play a role in endoplasmic reticulum (ER) and nuclear envelope dynamics, including assembly of the nuclear pore complex and maintenance of nuclear envelope structural integrity (73, 75). Torsin-1B is primarily localized to the ER and the perinuclear space (the space between the inner and outer nuclear membranes). It is also associated with the nuclear envelope where it is thought to interact with other nuclear and ER membrane proteins. Torsin-1B contains: *i*) an N-terminal domain involved in the regulation of its activity and possibly in substrate recognition; *ii*) an AAA+ ATPase domain responsible for ATP binding and hydrolysis; and *iii*) a C-terminal domain involved in protein-protein interactions and possibly in anchoring the protein to the nuclear envelope (76). Key amino acid sites include Glu178, which is essential for ATPase activity (73, 77). Based on the AlphaFold prediction, Torsin-1B adopts the typical AAA+ ATPase fold, which includes a large alpha/beta domain for ATP binding and a smaller domain involved in substrate interaction and oligomerization.

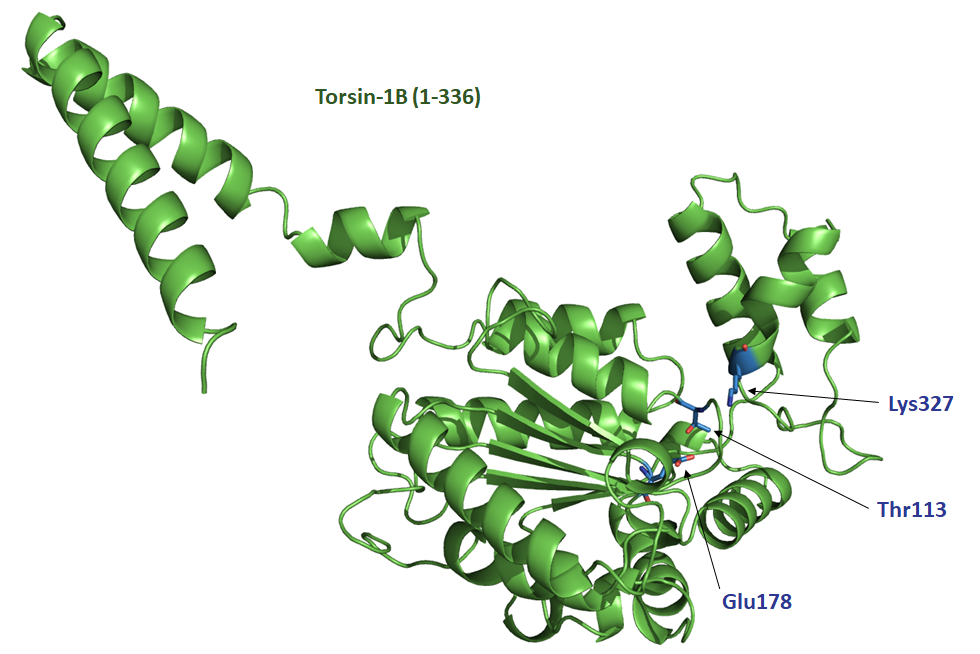

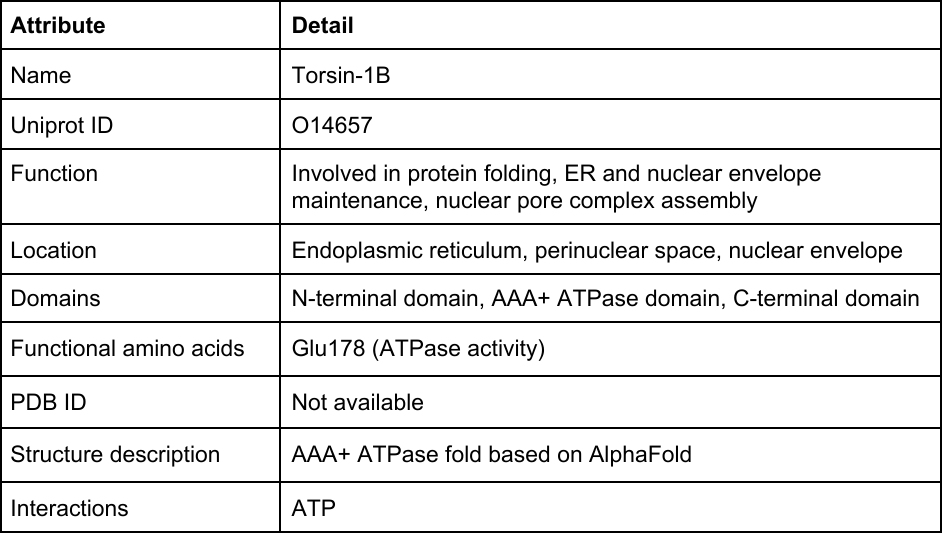

**YddG (Uniprot: D7A5Q8, PDB ID: 5I20)**

The protein YddG is a member of the major facilitator superfamily (MFS) of transporters. YddG is involved in the efflux of aromatic amino acids and their derivatives, such as tryptophan, phenylalanine and tyrosine, from the cell (78). This efflux activity is critical for maintaining intracellular amino acid levels and preventing the accumulation of potentially toxic compounds (78, 79). YddG plays an important role in bacterial physiology, particularly under stress conditions where the regulation of intracellular metabolites is essential for survival. YddG is an integral membrane protein located in the inner membrane of *Escherichia coli*. Its membrane localization allows it to function effectively as a transporter, facilitating the movement of substrates across the lipid bilayer (79). YddG consists of 10 transmembrane helices that form the transport channel across the membrane (79). The intracellular and extracellular loops connect the transmembrane helices and are involved in substrate recognition and conformational changes of the transporter, while the N- and C-terminal regions are important for the overall stability and function of the transporter. Specific amino acids within the transmembrane helices and loops are critical for YddG function. Such amino acid sites include Tyr78, HIs79, Tyr82, Trp101 and Trp163, all of which are essential for threonine and/or methionine uptake (79). The three-dimensional structure of YddG adopts the typical MFS fold, characterized by a large central cavity formed by the transmembrane helices. This cavity undergoes conformational changes during the transport cycle, alternating between inward-facing and outward-facing states to facilitate substrate movement across the membrane (79). YddG functions via a proton-motive force-dependent mechanism in which the energy derived from the proton gradient across the membrane drives the efflux of aromatic amino acids.

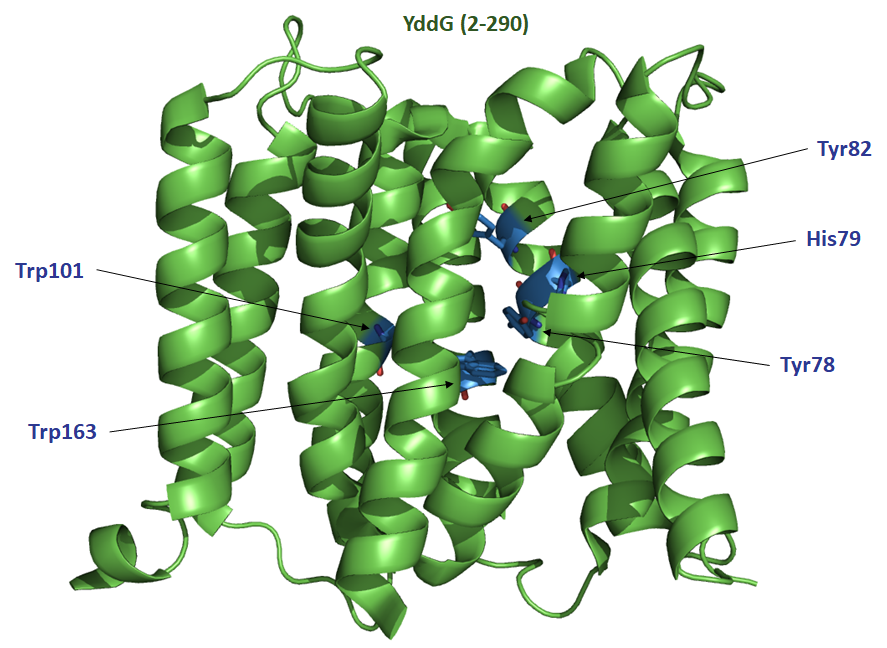

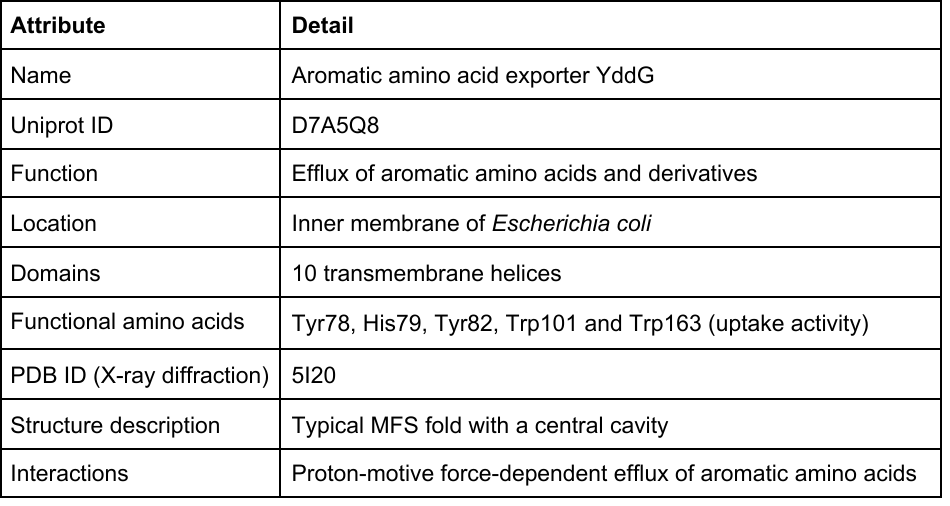

**GDP-mannose transporter 1 (Uniprot: P40107, PDB ID: 5OGE)**

GDP-mannose transporter 1, encoded by the VRG4 gene, is essential for transporting GDP-mannose from the cytosol into the lumen of the Golgi apparatus (80, 81). GDP-mannose is a critical substrate for glycosylation processes, including protein glycosylation and the synthesis of glycosylphosphatidylinositol (GPI) anchors (82, 83). By supplying GDP-mannose to the Golgi lumen, VRG4 plays a vital role in the proper functioning of the secretory pathway and cell wall biogenesis in yeast. VRG4 is localized in the Golgi membrane, where it functions as a transmembrane transporter (84). Its positioning in the Golgi apparatus is essential for its role in delivering GDP-mannose to the glycosylation machinery (85). VRG4 consists of 10 transmembrane helices that form the transport channel across the membrane (84). The intracellular and extracellular loops connect the transmembrane helices and are involved in substrate recognition and transport regulation. Some amino acid sites were reported to be essential in transport activities, such as Tyr28, Tyr114, Tyr281 and Lys289 (84). As YddG, VRG4 is a member of the major facilitator superfamily (MFS) of transporters, and adopts the typical MFS fold, characterized by a large central cavity formed by the transmembrane helices (84). VRG4 functions through an antiport mechanism, where the transport of GDP-mannose into the Golgi lumen is coupled with the export of GMP (guanosine monophosphate) to the cytosol (81). This exchange is driven by the concentration gradients of the substrates, ensuring efficient delivery of GDP-mannose for glycosylation processes.

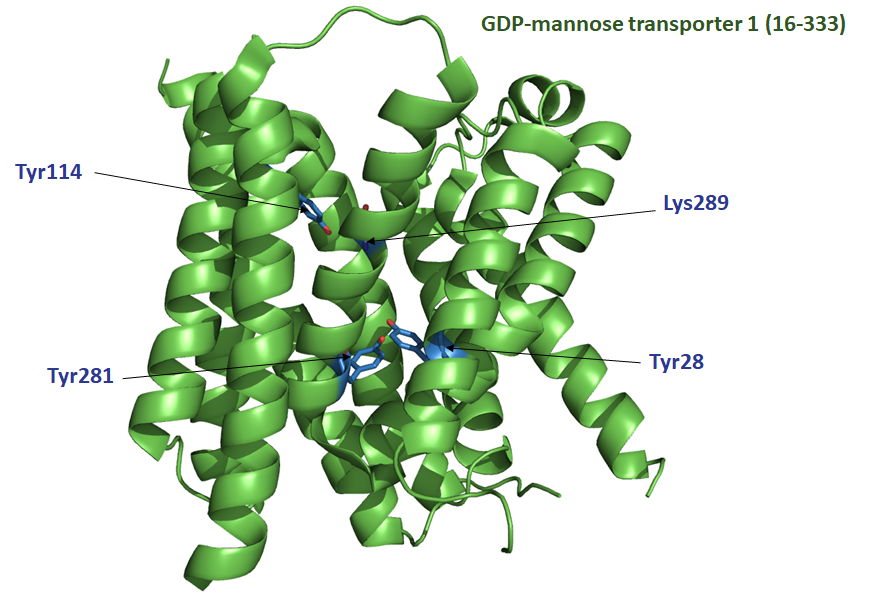

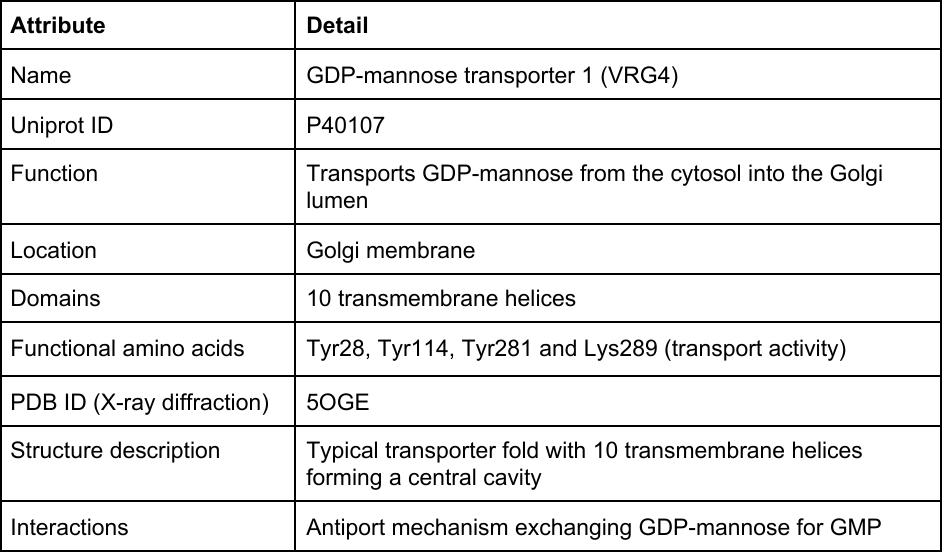

**GTPase HRas (Uniprot: P01112, PDB ID: 5P21)**

HRas is a small GTPase that belongs to the Ras superfamily of proteins, which play a critical role in cellular signaling. HRas functions as a molecular switch, cycling between an active GTP-bound state and an inactive GDP-bound state (86, 87). It is involved in the transduction of signals from cell surface receptors to intracellular signaling pathways, thereby influencing various cellular processes such as proliferation, differentiation and survival. HRas is particularly important in the Ras/MAPK signaling pathway, which is critical for cell growth and development (86, 87). HRas is primarily located at the inner surface of the plasma membrane where it associates with lipid rafts. It can also be found in the cytosol when not associated with membranes. Membrane association is facilitated by post-translational modifications, including farnesylation, which anchor HRas to the membrane (88, 89). HRas is a globular protein of 189 amino acids consisting of: *i*) a GTPase domain responsible for binding and hydrolyzing GTP; and *ii*) a C-terminal hypervariable region containing the sites for post-translational modifications that undergo farnesylation, proteolysis, and methylation, which are critical for membrane association (90). The amino acids Glu31, Asp33, Pro34, Ile36, Glu37, Asp38, Ser39, Tyr40, and Arg41 have been reported to be important for the enhancement of GTPase activity by GTPase-activating proteins (GAPs) (91–94). The three-dimensional structure of HRas reveals a central beta-sheet surrounded by alpha-helices, typical of the GTPase fold (PDB ID: 5P21) (90). HRas is activated when GTP replaces GDP, which is facilitated by guanine nucleotide exchange factors (GEFs) (86, 87). In the GTP-bound state, HRas interacts with downstream effectors such as RAF kinase and initiates signaling cascades such as the MAPK/ERK pathway. GAPs enhance the intrinsic GTPase activity of HRas, leading to the hydrolysis of GTP to GDP and returning HRas to its inactive state.

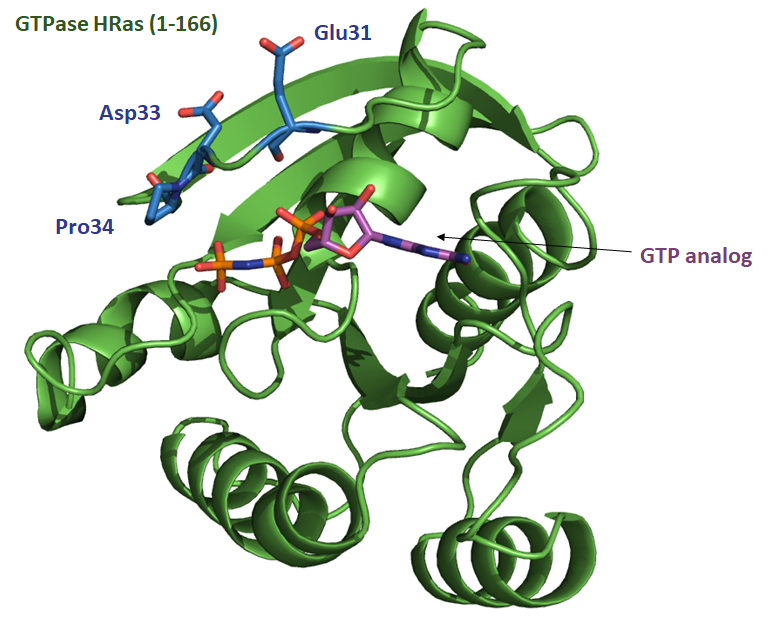

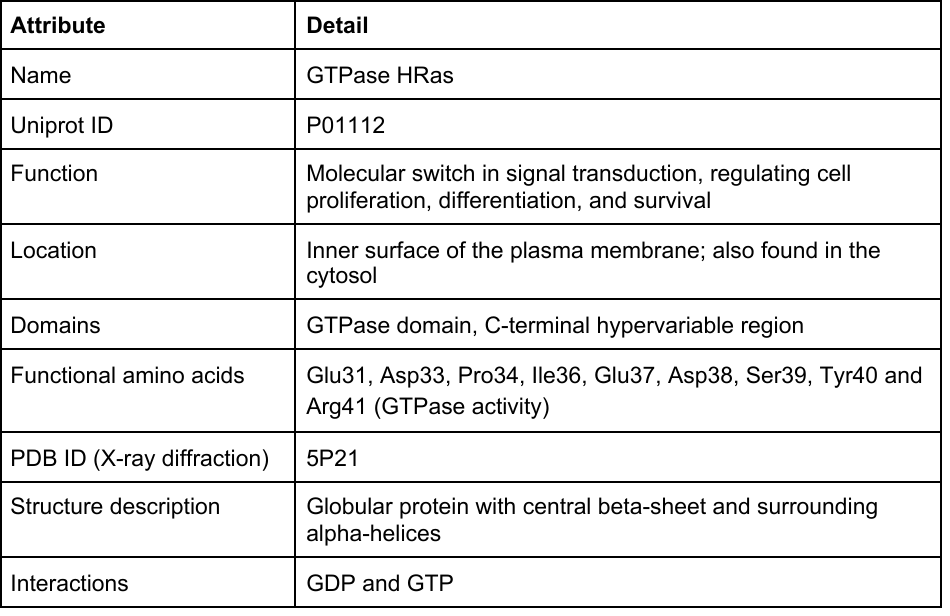

**Supplementary Data S2 – Detailed results of CONSTRUCT for other case studies**

**Cytochrome c (Uniprot: P99999, PDB ID: 1J3S)**

Rate4Site was first run on a dataset of 853 Cytochrome c orthologous sequences. The most conserved amino acid sites were uniformly distributed throughout the tertiary structure of Cytochrome c (*left structure*). Of note, these conserved sites included Lys13, which is located in the heme binding site (95). CONSTRUCT was then run on the same dataset. A patch of conserved amino acid sites was detected (log(*p*-value) = 15.99) at an optimal distance of 13 Å. This patch covered the heme binding site (*right structure*). The patch contained Lys13, but also Gln16, His18, Lys79 and Ile81, which coordinate the heme iron and are important for the electron transfer function (6–8, 95).

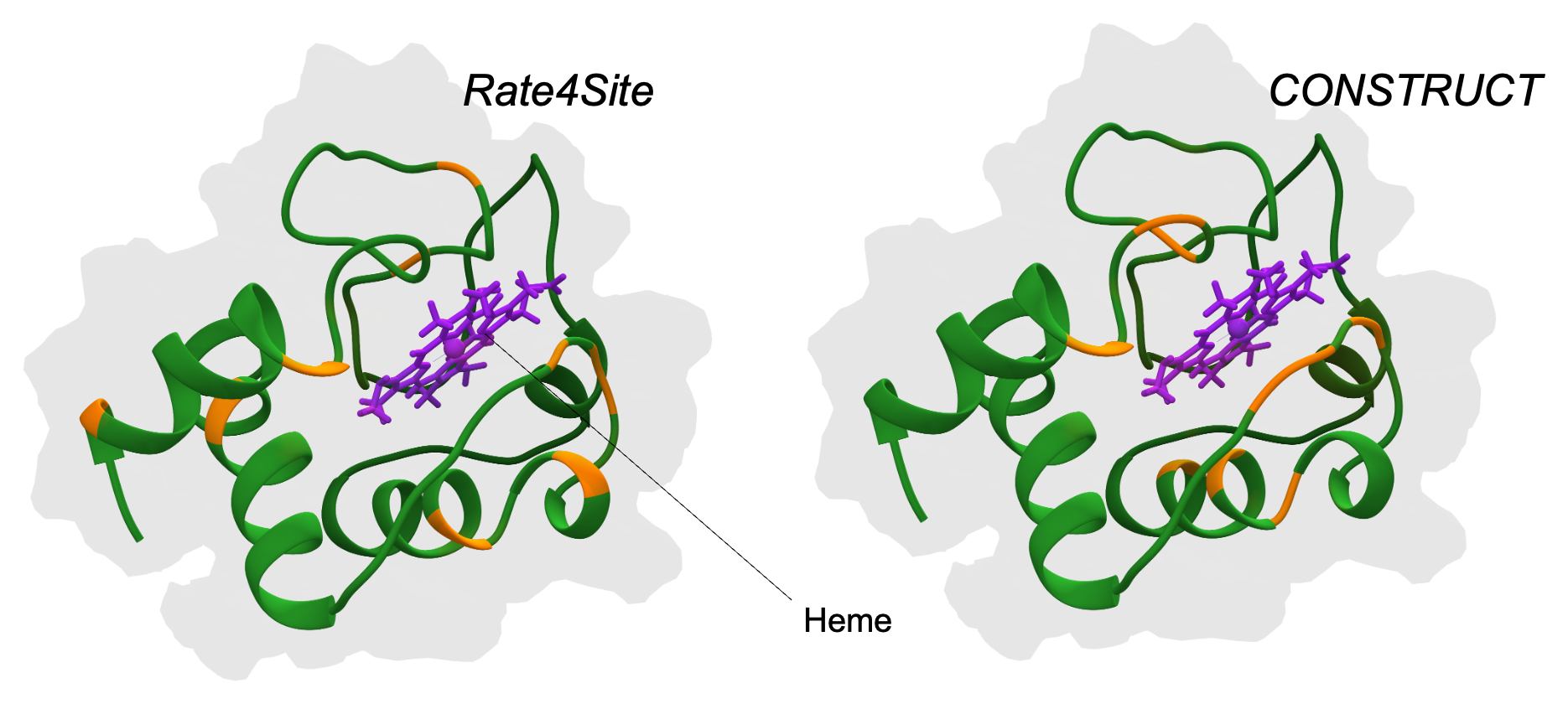

**DHFR (Uniprot: A7UD81, PDB ID: 3QGT)**

Rate4Site was first run on a dataset of 366 DHFR orthologous sequences. Again, the most conserved amino acid sites were uniformly distributed throughout the tertiary structure (*left structure*). These conserved sites included Ala16, which is associated with pyrimethamine resistance when mutated to valine and is located in the binding pocket with NADPH or pyrimethamine (27, 96). CONSTRUCT was then run on the same dataset. A patch of conserved amino acid sites was detected (log(*p*-value) = 72.82) at an optimal distance of 16 Å. This patch covered the binding site with NADPH or antimalarial drugs such as pyrimethamine (*right structure*). Especially, the patch included Ser108 and Ile112, both of which are located in the binding pocket and are involved in NADPH/pyrimethamine interaction (28, 96).

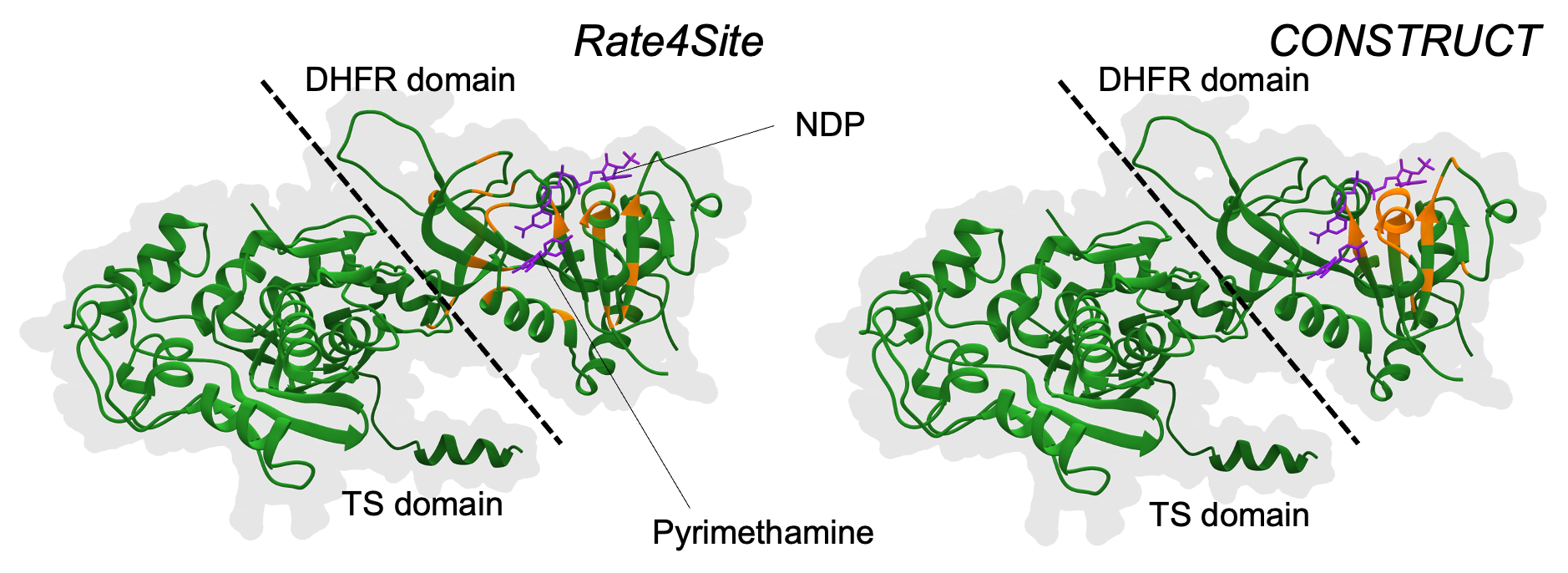

**Myoglobin (Uniprot: P02144, PDB ID: 3RGK)**

Using the Rate4Site algorithm, which ignores the spatial correlation of site-specific substitution rates in protein tertiary structure, conserved amino acid sites were widely distributed throughout the protein structure based on 454 orthologous sequences (*left structure*). Using CONSTRUCT, a spatial correlation of site-specific substitution rates was detected: the maximum strength of spatial correlation was observed at a distance of 20 Å, associated with a log(*p*-value) of 41.86. The most conserved sites identified by CONSTRUCT formed a well-defined patch in the tertiary structure (*right structure*). In addition, the conserved patch of amino acid sites overlapped with the binding site of binuclear Cu(II) for hydrolytic cleavage of the protein (33). The patch included the amino acid sites Gln91, Ser92, Ala94 and Thr95, all of which are cleaved by Cu(II) (33).

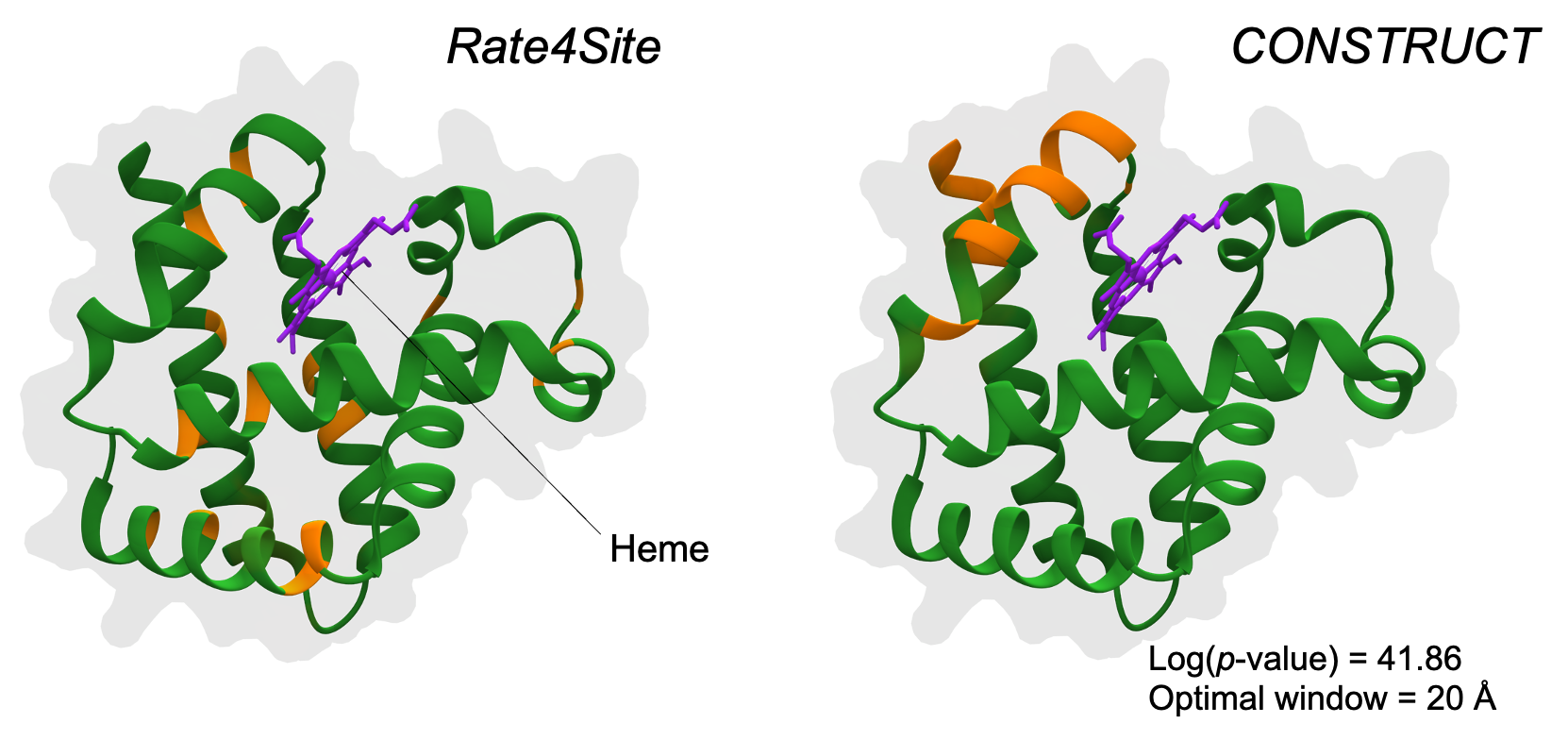

**DHPS (Uniprot: Q25704, PDB ID: 6JWV)**

The DHPS dataset consisted of 55 orthologous sequences. Rate4site revealed that the most conserved sites were largely distributed throughout the protein tertiary structure (*left structure*). However, these sites included Ser436 and Lys609, which are involved in pteroate and sulfa derivatives (46). CONSTRUCT was then applied, and the maximum strength of spatial correlation was detected at a distance of 19 Å, associated with a log(*p*-value) of 60.62. Again, the most conserved sites identified by CONSTRUCT formed a well-defined patch in the tertiary structure previously reported as the site of sulfa interaction (*right structure*). In particular, the patch included the amino acid sites Asn502, Asp539, Phe580, Lys609, Arg686 and His699, which are important for pteroate and/or sulfa derivative interaction (46).

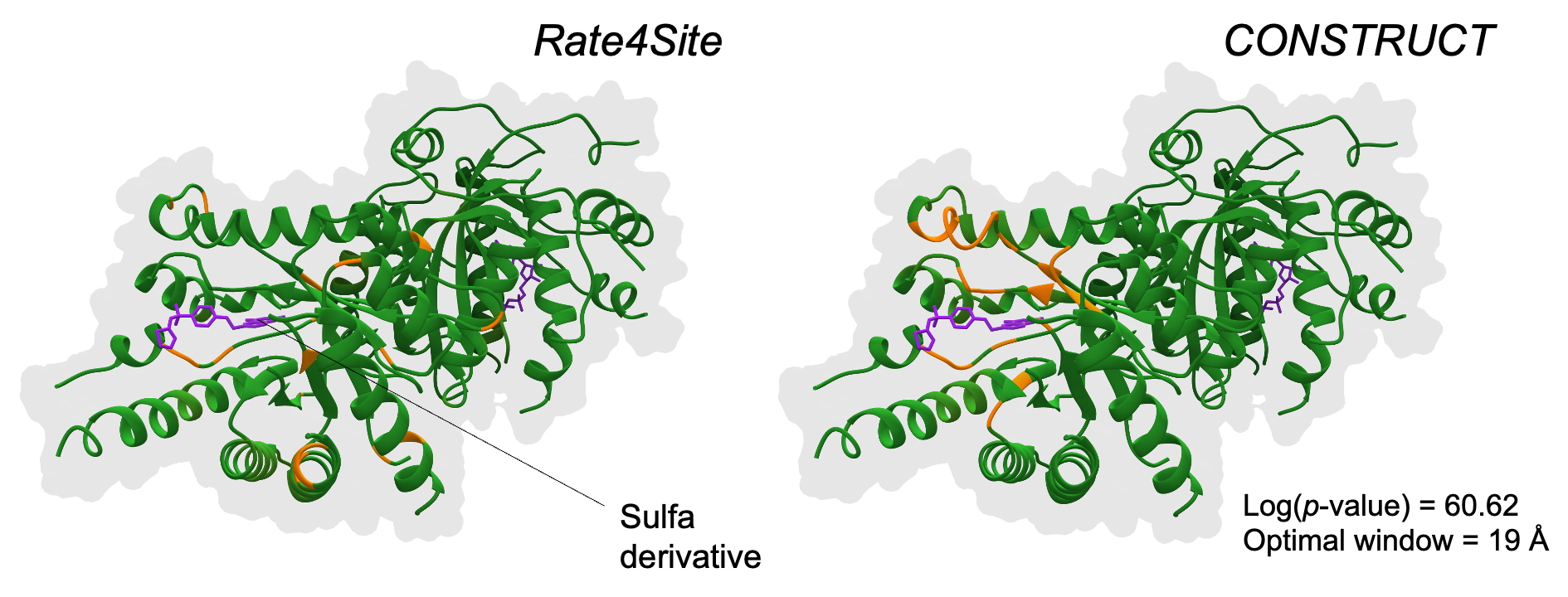

**MAPK1 (Uniprot: P63086, PDB ID: 5UMO)**

Rate4Site was first run on a dataset of 493 MAPK1 orthologous sequences. The most conserved amino acid sites were uniformly distributed throughout the tertiary structure (*left structure*). However, these conserved sites included Asp147, which is part of the catalytic loop and essential for ATP binding and catalysis (66). CONSTRUCT was then run on the same dataset. A patch of conserved amino acid sites was detected (log(*p*-value) = 50.73) at an optimal distance of 17 Å. This patch covered the catalytic site of the protein (*right structure*) (66). Especially, the patch included Arg146, Tyr185, Val186, Ala187, and Ser211, which are both phosphorylated sites and interacting sites with ATP (66).

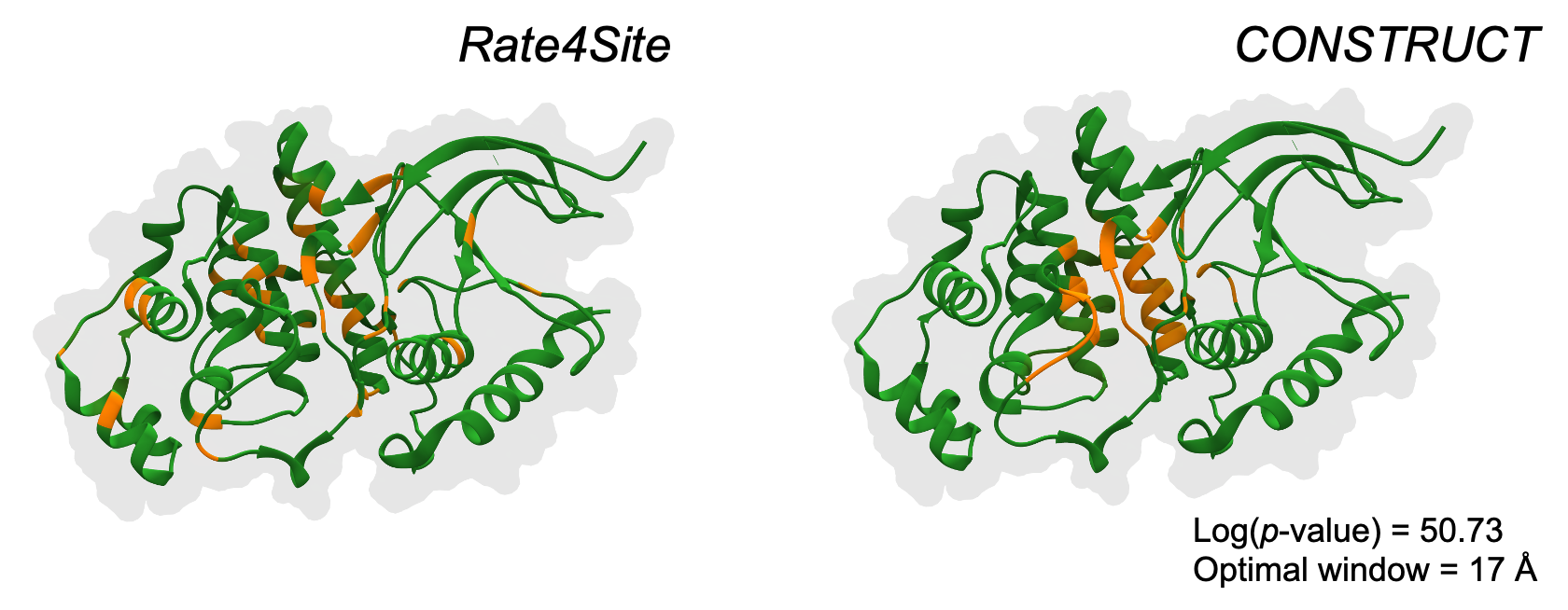

**SGLT1 (Uniprot: P13866, PDB ID: 7SL8)**

Rate4Site was first run on a dataset of 402 SGLT1 orthologous sequences. The most conserved amino acid sites were uniformly distributed throughout the tertiary structure (*left structure*). However, these conserved sites included Glu102 and Lys321, which are located in the glucose binding pocket (72). CONSTRUCT was then run on the same dataset. A patch of conserved amino acid sites was detected (log(*p*-value) = 88.76) at an optimal distance of 20 Å. This patch covered both the extracellular gate of the protein and the glucose binding pocket (*right structure*) (72). Especially, we found amino acid sites Asn78 and Tyr290, which have been shown to be involved in glucose binding; and Gly86, Leu452, Phe453, which are located at the extracellular gate of SGLT1 (72).

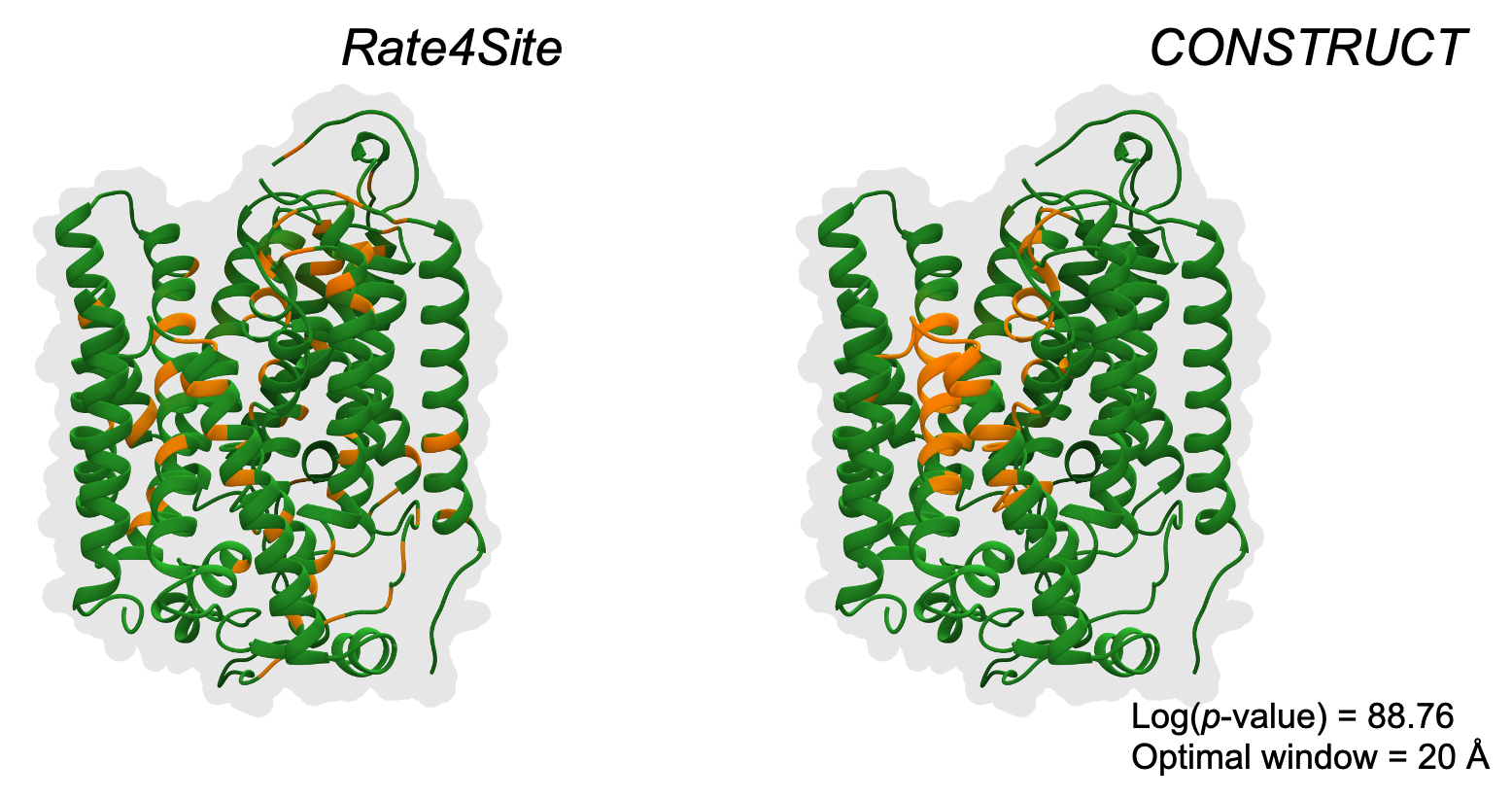

**GTPase HRas (Uniprot: P01112, PDB ID: 5P21)**

The HRas dataset consisted of 421 orthologous sequences. Rate4site revealed that the most conserved sites were largely distributed throughout the protein tertiary structure (*left structure*). However, these sites included Phe28, Asn116, Asp119, Thr144 and Ser145, which are involved in the interaction with GppNp molecule (90). CONSTRUCT was then applied, and the maximum strength of spatial correlation was detected at a distance of 17 Å, associated with a log(*p*-value) of 29.44. The most conserved sites identified by CONSTRUCT formed a well-defined patch in the tertiary structure previously reported as the interaction site between RAS and its effectors (*right structure*) (90). In particular, the patch included the amino acid sites Lys16, Ser17, Glu31, Tyr32, Asp33, Pro34, Thr35, Glu37, Asp38, Asp57, Gly60, Glu62 and Glu63, all of which were previously reported to be involved in such interactions (90).

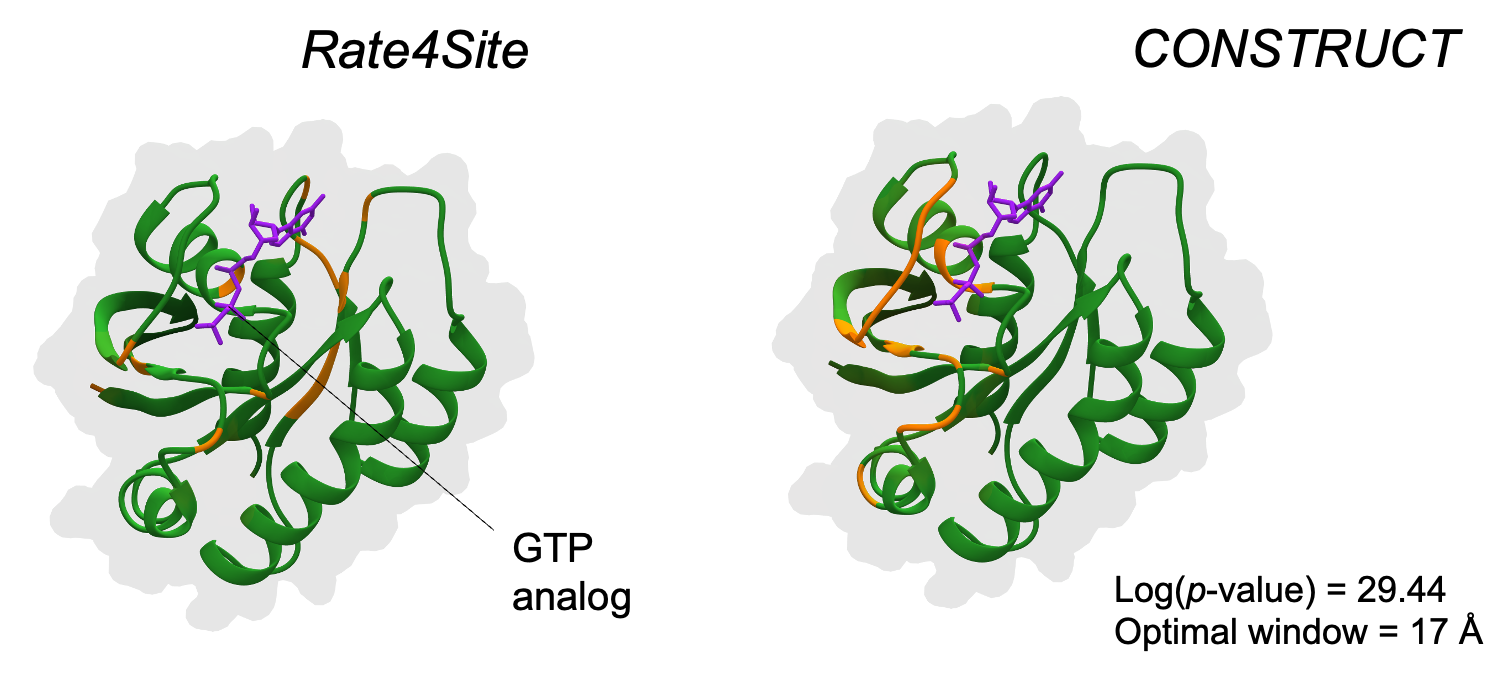

**Supplementary Table S1 – Analysis duration for each case study.**

| **Protein** | **Number of orthologous sequences** | **Size of the alignment** | **Analysis duration (s)** |
| --- | --- | --- | --- |
| Cytochrome c | 853 | 103 | 423 |
| MDM2 | 376 | 109 | 88 |
| KEAP1 (propeller) | 135 | 285 | 35 |
| DHFR (N-terminal domain) | 366 | 617 | 571 |
| Myoglobin | 454 | 153 | 180 |
| cAMP-dependent protein kinase A | 254 | 334 | 182 |
| DHPS (catalytic domain) | 55 | 641 | 24 |
| CFTR | 415 | 1,463 | 1,411 |
| MAPK1 | 493 | 365 | 701 |
| SGLT1 | 402 | 667 | 1,044 |
| Torsin-1B | 299 | 338 | 295 |
| YddG | 383 | 277 | 294 |
| GDP-mannose transporter 1 | 366 | 322 | 310 |
| GTPase Hras | 421 | 166 | 202 |

**Supplementary Table S2 – List of software and packages used in CONSTRUCT.**

| **Software** | | **Version** | **Link** |
| --- | --- | --- | --- |
|  | Rate4Site | 3.0.0 | https://www.tau.ac.il/~itaymay/cp/rate4site.html |
|  | Python | 3.10.12 | https://www.python.org/ |
|  | R | 4.1.2 | https://www.r-project.org/ |
| **Python packages** | |  |  |
|  | Tkinter | 8.6 | https://docs.python.org/fr/3/library/tkinter.html |
|  | Customtkinter | 0.3 | https://pypi.org/project/customtkinter/0.3/ |
| **R packages** | |  |  |
|  | Tidyverse | 2.0.0 | https://www.tidyverse.org/ |
|  | Bio3d | 2.4-4 | http://thegrantlab.org/bio3d/ |
|  | BiocManager | 1.30.23 | https://www.bioconductor.org/ |
|  | Msa | 1.26.0 | https://bioconductor.org/packages/release/bioc/html/msa.html |

Note – All the packages and software (except Python and R) can be easily installed using a bash script provided with CONSTRUCT.

**References**

1. Li,K., Li,Y., Shelton,J.M., Richardson,J.A., Spencer,E., Chen,Z.J., Wang,X. and Williams,R.S. (2000) Cytochrome c deficiency causes embryonic lethality and attenuates stress-induced apoptosis. *Cell*, **101**, 389–399.

2. Garrido,C., Galluzzi,L., Brunet,M., Puig,P.E., Didelot,C. and Kroemer,G. (2006) Mechanisms of cytochrome c release from mitochondria. *Cell Death Differ*, **13**, 1423–1433.

3. Zou,H., Li,Y., Liu,X. and Wang,X. (1999) An APAF-1.cytochrome c multimeric complex is a functional apoptosome that activates procaspase-9. *J Biol Chem*, **274**, 11549–11556.

4. Bernardi,P. and Azzone,G.F. (1981) Cytochrome c as an electron shuttle between the outer and inner mitochondrial membranes. *J Biol Chem*, **256**, 7187–7192.

5. Kang,X. and Carey,J. (1999) Role of heme in structural organization of cytochrome c probed by semisynthesis. *Biochemistry*, **38**, 15944–15951.

6. Bushnell,G.W., Louie,G.V. and Brayer,G.D. (1990) High-resolution three-dimensional structure of horse heart cytochrome c. *J Mol Biol*, **214**, 585–595.

7. Döpner,S., Hildebrandt,P., Rosell,F.I., Mauk,A.G., von Walter,M., Buse,G. and Soulimane,T. (1999) The structural and functional role of lysine residues in the binding domain of cytochrome c in the electron transfer to cytochrome c oxidase. *Eur J Biochem*, **261**, 379–391.

8. Moreno-Beltrán,B., Díaz-Moreno,I., González-Arzola,K., Guerra-Castellano,A., Velázquez-Campoy,A., De la Rosa,M.A. and Díaz-Quintana,A. (2015) Respiratory complexes III and IV can each bind two molecules of cytochrome c at low ionic strength. *FEBS Lett*, **589**, 476–483.

9. Kussie,P.H., Gorina,S., Marechal,V., Elenbaas,B., Moreau,J., Levine,A.J. and Pavletich,N.P. (1996) Structure of the MDM2 Oncoprotein Bound to the p53 Tumor Suppressor Transactivation Domain. *Science*, **274**, 948–953.

10. Momand,J., Wu,H.H. and Dasgupta,G. (2000) MDM2--master regulator of the p53 tumor suppressor protein. *Gene*, **242**, 15–29.

11. Roth,J., Dobbelstein,M., Freedman,D.A., Shenk,T. and Levine,A.J. (1998) Nucleo-cytoplasmic shuttling of the hdm2 oncoprotein regulates the levels of the p53 protein via a pathway used by the human immunodeficiency virus rev protein. *EMBO J*, **17**, 554–564.

12. Linke,K., Mace,P.D., Smith,C.A., Vaux,D.L., Silke,J. and Day,C.L. (2008) Structure of the MDM2/MDMX RING domain heterodimer reveals dimerization is required for their ubiquitylation in trans. *Cell Death Differ*, **15**, 841–848.

13. Lai,Z., Freedman,D.A., Levine,A.J. and McLendon,G.L. (1998) Metal and RNA Binding Properties of the hdm2 RING Finger Domain. *Biochemistry*, **37**, 17005–17015.

14. Moll,U.M. and Petrenko,O. (2003) The MDM2-p53 Interaction. *Molecular Cancer Research*, **1**, 1001–1008.

15. Weber,J.D., Taylor,L.J., Roussel,M.F., Sherr,C.J. and Bar-Sagi,D. (1999) Nucleolar Arf sequesters Mdm2 and activates p53. *Nat Cell Biol*, **1**, 20–26.

16. Zhang,D.D. and Hannink,M. (2003) Distinct Cysteine Residues in Keap1 Are Required for Keap1-Dependent Ubiquitination of Nrf2 and for Stabilization of Nrf2 by Chemopreventive Agents and Oxidative Stress. *Mol Cell Biol*, **23**, 8137–8151.

17. Zhang,D.D., Lo,S.-C., Cross,J.V., Templeton,D.J. and Hannink,M. (2004) Keap1 is a redox-regulated substrate adaptor protein for a Cul3-dependent ubiquitin ligase complex. *Mol Cell Biol*, **24**, 10941–10953.

18. Zhang,D.D., Lo,S.-C., Sun,Z., Habib,G.M., Lieberman,M.W. and Hannink,M. (2005) Ubiquitination of Keap1, a BTB-Kelch substrate adaptor protein for Cul3, targets Keap1 for degradation by a proteasome-independent pathway. *J Biol Chem*, **280**, 30091–30099.

19. Rachakonda,G., Xiong,Y., Sekhar,K.R., Stamer,S.L., Liebler,D.C. and Freeman,M.L. (2008) Covalent modification at Cys151 dissociates the electrophile sensor Keap1 from the ubiquitin ligase CUL3. *Chem Res Toxicol*, **21**, 705–710.

20. Canning,P., Cooper,C.D.O., Krojer,T., Murray,J.W., Pike,A.C.W., Chaikuad,A., Keates,T., Thangaratnarajah,C., Hojzan,V., Marsden,B.D., *et al.* (2013) Structural basis for Cul3 protein assembly with the BTB-Kelch family of E3 ubiquitin ligases. *J Biol Chem*, **288**, 7803–7814.

21. Canning,P., Sorrell,F.J. and Bullock,A.N. (2015) Structural basis of Keap1 interactions with Nrf2. *Free Radic Biol Med*, **88**, 101–107.

22. Lo,S., Li,X., Henzl,M.T., Beamer,L.J. and Hannink,M. (2006) Structure of the Keap1:Nrf2 interface provides mechanistic insight into Nrf2 signaling. *The EMBO Journal*, **25**, 3605–3617.

23. Rastelli,G., Sirawaraporn,W., Sompornpisut,P., Vilaivan,T., Kamchonwongpaisan,S., Quarrell,R., Lowe,G., Thebtaranonth,Y. and Yuthavong,Y. (2000) Interaction of pyrimethamine, cycloguanil, WR99210 and their analogues with Plasmodium falciparum dihydrofolate reductase: structural basis of antifolate resistance. *Bioorg Med Chem*, **8**, 1117–1128.

24. Dasgupta,T. and Anderson,K.S. (2008) Probing the Role of Parasite-specific, Distant, Structural Regions on Communication and Catalysis in the Bifunctional Thymidylate Synthase- Dihydrofolate Reductase from Plasmodium falciparum. *Biochemistry*, **47**, 1336–1345.

25. Sirawaraporn,W., Prapunwattana,P., Sirawaraporn,R., Yuthavong,Y. and Santi,D.V. (1993) The dihydrofolate reductase domain of Plasmodium falciparum thymidylate synthase-dihydrofolate reductase. Gene synthesis, expression, and anti-folate-resistant mutants. *Journal of Biological Chemistry*, **268**, 21637–21644.

26. Sirawaraporn,W., Sirawaraporn,R., Yongkiettrakul,S., Anuwatwora,A., Rastelli,G., Kamchonwongpaisan,S. and Yuthavong,Y. (2002) Mutational analysis of *Plasmodium falciparum* dihydrofolate reductase: the role of aspartate 54 and phenylalanine 223 on catalytic activity and antifolate binding. *Molecular and Biochemical Parasitology*, **121**, 185–193.

27. Yuthavong,Y. (2002) Basis for antifolate action and resistance in malaria. *Microbes and Infection*, **4**, 175–182.

28. Vanichtanankul,J., Taweechai,S., Yuvaniyama,J., Vilaivan,T., Chitnumsub,P., Kamchonwongpaisan,S. and Yuthavong,Y. (2011) Trypanosomal Dihydrofolate Reductase Reveals Natural Antifolate Resistance. *ACS Chem. Biol.*, **6**, 905–911.

29. Ordway,G.A. and Garry,D.J. (2004) Myoglobin: an essential hemoprotein in striated muscle. *J Exp Biol*, **207**, 3441–3446.

30. Garry,D.J. and Mammen,P.P.A. (2007) Molecular insights into the functional role of myoglobin. *Adv Exp Med Biol*, **618**, 181–193.

31. Braunlin,E.A., Wahler,G.M., Swayze,C.R., Lucas,R.V. and Fox,I.J. (1986) Myoglobin facilitated oxygen diffusion maintains mechanical function of mammalian cardiac muscle. *Cardiovasc Res*, **20**, 627–636.

32. Vojtechovský,J., Chu,K., Berendzen,J., Sweet,R.M. and Schlichting,I. (1999) Crystal structures of myoglobin-ligand complexes at near-atomic resolution. *Biophys J*, **77**, 2153–2174.

33. Zhang,L., Mei,Y., Zhang,Y., Li,S., Sun,X. and Zhu,L. (2003) Regioselective cleavage of myoglobin with copper(II) compounds at neutral pH. *Inorg Chem*, **42**, 492–498.

34. Enyenihi,A.A., Yang,H., Ytterberg,A.J., Lyutvinskiy,Y. and Zubarev,R.A. (2011) Heme Binding in Gas-Phase Holo-Myoglobin Cations: Distal Becomes Proximal? *J. Am. Soc. Mass Spectrom.*, **22**, 1763–1770.

35. Hubbard,S.R., Hendrickson,W.A., Lambright,D.G. and Boxer,S.G. (1990) X-ray crystal structure of a recombinant human myoglobin mutant at 2·8 Å resolution. *Journal of Molecular Biology*, **213**, 215–218.

36. Turnham,R.E. and Scott,J.D. (2016) Protein kinase A catalytic subunit isoform PRKACA; History, function and physiology. *Gene*, **577**, 101–108.

37. Nakashima,S. (2002) Protein Kinase Cα (PKCα): Regulation and Biological Function. *The Journal of Biochemistry*, **132**, 669–675.

38. Cheung,J., Ginter,C., Cassidy,M., Franklin,M.C., Rudolph,M.J., Robine,N., Darnell,R.B. and Hendrickson,W.A. (2015) Structural insights into mis-regulation of protein kinase A in human tumors. *Proceedings of the National Academy of Sciences*, **112**, 1374–1379.

39. Taylor,S.S., Søberg,K., Kobori,E., Wu,J., Pautz,S., Herberg,F.W. and Skålhegg,B.S. (2022) The Tails of Protein Kinase A. *Mol Pharmacol*, **101**, 219–225.

40. Toyota,A., Goto,M., Miyamoto,M., Nagashima,Y., Iwasaki,S., Komatsu,T., Momose,T., Yoshida,K., Tsukada,T., Matsufuji,T., *et al.* (2022) Novel protein kinase cAMP-Activated Catalytic Subunit Alpha (PRKACA) inhibitor shows anti-tumor activity in a fibrolamellar hepatocellular carcinoma model. *Biochem Biophys Res Commun*, **621**, 157–161.

41. Yang,J., Ten Eyck,L.F., Xuong,N.-H. and Taylor,S.S. (2004) Crystal Structure of a cAMP-dependent Protein Kinase Mutant at 1.26   Å: New Insights into the Catalytic Mechanism. *Journal of Molecular Biology*, **336**, 473–487.

42. Søberg,K. and Skålhegg,B.S. (2018) The Molecular Basis for Specificity at the Level of the Protein Kinase a Catalytic Subunit. *Front Endocrinol (Lausanne)*, **9**, 538.

43. Tomasini,M.D., Wang,Y., Karamafrooz,A., Li,G., Beuming,T., Gao,J., Taylor,S.S., Veglia,G. and Simon,S.M. (2018) Conformational Landscape of the PRKACA-DNAJB1 Chimeric Kinase, the Driver for Fibrolamellar Hepatocellular Carcinoma. *Sci Rep*, **8**, 720.

44. Then,R. (2007) Dihydropteroate Synthase*. In Enna,S.J., Bylund,D.B. (eds), *xPharm: The Comprehensive Pharmacology Reference*. Elsevier, New York, pp. 1–7.

45. Hyde,J.E. (2005) Exploring the folate pathway in Plasmodium falciparum. *Acta Trop*, **94**, 191–206.

46. Chitnumsub,P., Jaruwat,A., Talawanich,Y., Noytanom,K., Liwnaree,B., Poen,S. and Yuthavong,Y. (2020) The structure of Plasmodium falciparum hydroxymethyldihydropterin pyrophosphokinase-dihydropteroate synthase reveals the basis of sulfa resistance. *The FEBS Journal*, **287**, 3273–3297.

47. Triglia,T. and Cowman,A.F. (1994) Primary structure and expression of the dihydropteroate synthetase gene of Plasmodium falciparum. *Proc Natl Acad Sci U S A*, **91**, 7149–7153.

48. Triglia,T., Menting,J.G.T., Wilson,C. and Cowman,A.F. (1997) Mutations in dihydropteroate synthase are responsible for sulfone and sulfonamide resistance in Plasmodium falciparum. *Proc Natl Acad Sci U S A*, **94**, 13944–13949.

49. Triglia,T., Wang,P., Sims,P.F., Hyde,J.E. and Cowman,A.F. (1998) Allelic exchange at the endogenous genomic locus in Plasmodium falciparum proves the role of dihydropteroate synthase in sulfadoxine-resistant malaria. *EMBO J*, **17**, 3807–3815.

50. Brooks,D.R., Wang,P., Read,M., Watkins,W.M., Sims,P.F. and Hyde,J.E. (1994) Sequence variation of the hydroxymethyldihydropterin pyrophosphokinase: dihydropteroate synthase gene in lines of the human malaria parasite, Plasmodium falciparum, with differing resistance to sulfadoxine. *Eur J Biochem*, **224**, 397–405.

51. Sheppard,D.N. and Welsh,M.J. (1999) Structure and function of the CFTR chloride channel. *Physiol Rev*, **79**, S23-45.

52. Higgins,C.F. (1992) Cystic fibrosis transmembrane conductance regulator (CFTR). *Br Med Bull*, **48**, 754–765.

53. Li,C., Ramjeesingh,M., Wang,W., Garami,E., Hewryk,M., Lee,D., Rommens,J.M., Galley,K. and Bear,C.E. (1996) ATPase activity of the cystic fibrosis transmembrane conductance regulator. *J Biol Chem*, **271**, 28463–28468.

54. Dörk,T., Wulbrand,U., Richter,T., Neumann,T., Wolfes,H., Wulf,B., Maass,G. and Tümmler,B. (1991) Cystic fibrosis with three mutations in the cystic fibrosis transmembrane conductance regulator gene. *Hum Genet*, **87**, 441–446.

55. Bobadilla,J.L., Macek,M., Fine,J.P. and Farrell,P.M. (2002) Cystic fibrosis: a worldwide analysis of CFTR mutations--correlation with incidence data and application to screening. *Hum Mutat*, **19**, 575–606.

56. Rowntree,R.K. and Harris,A. (2003) The phenotypic consequences of CFTR mutations. *Ann Hum Genet*, **67**, 471–485.

57. Devidas,S. and Guggino,W.B. (1997) CFTR: domains, structure, and function. *J Bioenerg Biomembr*, **29**, 443–451.

58. Powe,A.C., Al-Nakkash,L., Li,M. and Hwang,T.-C. (2002) Mutation of Walker-A lysine 464 in cystic fibrosis transmembrane conductance regulator reveals functional interaction between its nucleotide-binding domains. *J Physiol*, **539**, 333–346.

59. Stratford,F.L.L., Ramjeesingh,M., Cheung,J.C., Huan,L.-J. and Bear,C.E. (2007) The Walker B motif of the second nucleotide-binding domain (NBD2) of CFTR plays a key role in ATPase activity by the NBD1-NBD2 heterodimer. *Biochem J*, **401**, 581–586.

60. Szollosi,A., Muallem,D.R., Csanády,L. and Vergani,P. (2011) Mutant cycles at CFTR’s non-canonical ATP-binding site support little interface separation during gating. *J Gen Physiol*, **137**, 549–562.

61. Lewis,H.A., Zhao,X., Wang,C., Sauder,J.M., Rooney,I., Noland,B.W., Lorimer,D., Kearins,M.C., Conners,K., Condon,B., *et al.* (2005) Impact of the ΔF508 Mutation in First Nucleotide-binding Domain of Human Cystic Fibrosis Transmembrane Conductance Regulator on Domain Folding and Structure*. *Journal of Biological Chemistry*, **280**, 1346–1353.

62. Chen,R.E. and Thorner,J. (2007) Function and regulation in MAPK signaling pathways: lessons learned from the yeast Saccharomyces cerevisiae. *Biochim Biophys Acta*, **1773**, 1311–1340.

63. Cargnello,M. and Roux,P.P. (2011) Activation and function of the MAPKs and their substrates, the MAPK-activated protein kinases. *Microbiol Mol Biol Rev*, **75**, 50–83.

64. Sugiura,R., Toda,T., Dhut,S., Shuntoh,H. and Kuno,T. (1999) The MAPK kinase Pek1 acts as a phosphorylation-dependent molecular switch. *Nature*, **399**, 479–483.

65. Whitehurst,A.W., Wilsbacher,J.L., You,Y., Luby-Phelps,K., Moore,M.S. and Cobb,M.H. (2002) ERK2 enters the nucleus by a carrier-independent mechanism. *Proc Natl Acad Sci U S A*, **99**, 7496–7501.

66. Zhang,F., Strand,A., Robbins,D., Cobb,M.H. and Goldsmith,E.J. (1994) Atomic structure of the MAP kinase ERK2 at 2.3 A resolution. *Nature*, **367**, 704–711.

67. Canagarajah,B.J., Khokhlatchev,A., Cobb,M.H. and Goldsmith,E.J. (1997) Activation Mechanism of the MAP Kinase ERK2 by Dual Phosphorylation. *Cell*, **90**, 859–869.

68. Turjanski,A.G., Hummer,G. and Gutkind,J.S. (2009) How Mitogen-Activated Protein Kinases Recognize and Phosphorylate Their Targets: A QM/MM Study. *J. Am. Chem. Soc.*, **131**, 6141–6148.

69. Martín,M.G., Turk,E., Lostao,M.P., Kerner,C. and Wright,E.M. (1996) Defects in Na+/glucose cotransporter (SGLT1) trafficking and function cause glucose-galactose malabsorption. *Nat Genet*, **12**, 216–220.

70. Hummel,C.S., Lu,C., Loo,D.D.F., Hirayama,B.A., Voss,A.A. and Wright,E.M. (2011) Glucose transport by human renal Na+/D-glucose cotransporters SGLT1 and SGLT2. *Am J Physiol Cell Physiol*, **300**, C14-21.

71. Erokhova,L., Horner,A., Ollinger,N., Siligan,C. and Pohl,P. (2016) The Sodium Glucose Cotransporter SGLT1 Is an Extremely Efficient Facilitator of Passive Water Transport. *J Biol Chem*, **291**, 9712–9720.

72. Han,L., Qu,Q., Aydin,D., Panova,O., Robertson,M.J., Xu,Y., Dror,R.O., Skiniotis,G. and Feng,L. (2022) Structure and mechanism of the SGLT family of glucose transporters. *Nature*, **601**, 274–279.

73. Rose,A.E., Zhao,C., Turner,E.M., Steyer,A.M. and Schlieker,C. (2014) Arresting a Torsin ATPase reshapes the endoplasmic reticulum. *J Biol Chem*, **289**, 552–564.

74. Hanson,P.I. and Whiteheart,S.W. (2005) AAA+ proteins: have engine, will work. *Nat Rev Mol Cell Biol*, **6**, 519–529.

75. Hewett,J.W., Kamm,C., Boston,H., Beauchamp,R., Naismith,T., Ozelius,L., Hanson,P.I., Breakefield,X.O. and Ramesh,V. (2004) TorsinB--perinuclear location and association with torsinA. *J Neurochem*, **89**, 1186–1194.

76. Laudermilch,E. and Schlieker,C. (2016) Torsin ATPases: structural insights and functional perspectives. *Curr Opin Cell Biol*, **40**, 1–7.

77. Jungwirth,M., Dear,M.L., Brown,P., Holbrook,K. and Goodchild,R. (2010) Relative tissue expression of homologous torsinB correlates with the neuronal specific importance of DYT1 dystonia-associated torsinA. *Hum Mol Genet*, **19**, 888–900.

78. Doroshenko,V., Airich,L., Vitushkina,M., Kolokolova,A., Livshits,V. and Mashko,S. (2007) YddG from Escherichia coli promotes export of aromatic amino acids. *FEMS Microbiol Lett*, **275**, 312–318.

79. Tsuchiya,H., Doki,S., Takemoto,M., Ikuta,T., Higuchi,T., Fukui,K., Usuda,Y., Tabuchi,E., Nagatoishi,S., Tsumoto,K., *et al.* (2016) Structural basis for amino acid export by DMT superfamily transporter YddG. *Nature*, **534**, 417–420.

80. Gao,X.D., Nishikawa,A. and Dean,N. (2001) Identification of a conserved motif in the yeast golgi GDP-mannose transporter required for binding to nucleotide sugar. *J Biol Chem*, **276**, 4424–4432.

81. Abe,M., Hashimoto,H. and Yoda,K. (1999) Molecular characterization of Vig4/Vrg4 GDP-mannose transporter of the yeast Saccharomyces cerevisiae. *FEBS Lett*, **458**, 309–312.

82. Ballou,L., Hitzeman,R.A., Lewis,M.S. and Ballou,C.E. (1991) Vanadate-resistant yeast mutants are defective in protein glycosylation. *Proc Natl Acad Sci U S A*, **88**, 3209–3212.

83. Kanik-Ennulat,C. and Neff,N. (1990) Vanadate-resistant mutants of Saccharomyces cerevisiae show alterations in protein phosphorylation and growth control. *Mol Cell Biol*, **10**, 898–909.

84. Parker,J.L. and Newstead,S. (2017) Structural basis of nucleotide sugar transport across the Golgi membrane. *Nature*, **551**, 521–524.

85. Hashimoto,H., Abe,M., Hirata,A., Noda,Y., Adachi,H. and Yoda,K. (2002) Progression of the stacked Golgi compartments in the yeast Saccharomyces cerevisiae by overproduction of GDP-mannose transporter. *Yeast*, **19**, 1413–1424.

86. Lander,H.M., Hajjar,D.P., Hempstead,B.L., Mirza,U.A., Chait,B.T., Campbell,S. and Quilliam,L.A. (1997) A molecular redox switch on p21(ras). Structural basis for the nitric oxide-p21(ras) interaction. *J Biol Chem*, **272**, 4323–4326.

87. Williams,J.G., Pappu,K. and Campbell,S.L. (2003) Structural and biochemical studies of p21Ras S-nitrosylation and nitric oxide-mediated guanine nucleotide exchange. *Proc Natl Acad Sci U S A*, **100**, 6376–6381.

88. Baker,R., Wilkerson,E.M., Sumita,K., Isom,D.G., Sasaki,A.T., Dohlman,H.G. and Campbell,S.L. (2013) Differences in the Regulation of K-Ras and H-Ras Isoforms by Monoubiquitination. *J Biol Chem*, **288**, 36856–36862.

89. Odeniyide,P., Yohe,M.E., Pollard,K., Vaseva,A.V., Calizo,A., Zhang,L., Rodriguez,F.J., Gross,J.M., Allen,A.N., Wan,X., *et al.* (2022) Targeting farnesylation as a novel therapeutic approach in HRAS-mutant rhabdomyosarcoma. *Oncogene*, **41**, 2973–2983.

90. Pai,E.F., Krengel,U., Petsko,G.A., Goody,R.S., Kabsch,W. and Wittinghofer,A. (1990) Refined crystal structure of the triphosphate conformation of H‐ras p21 at 1.35 A resolution: implications for the mechanism of GTP hydrolysis. *The EMBO Journal*, **9**, 2351–2359.

91. Nyíri,K., Koppány,G. and Vértessy,B.G. (2020) Structure-based inhibitor design of mutant RAS proteins—a paradigm shift. *Cancer Metastasis Rev*, **39**, 1091–1105.

92. Bueno,A., Morilla,I., Diez,D., Moya-Garcia,A.A., Lozano,J. and Ranea,J.A.G. (2016) Exploring the interactions of the RAS family in the human protein network and their potential implications in RAS-directed therapies. *Oncotarget*, **7**, 75810–75826.

93. Lu,J., Bera,A.K., Gondi,S. and Westover,K.D. (2018) KRAS Switch Mutants D33E and A59G Crystallize in the State 1 Conformation. *Biochemistry*, **57**, 324–333.

94. Kumar,S., Sharma,D., Narasimhan,B., Ramasamy,K., Shah,S.A.A., Lim,S.M. and Mani,V. (2019) Computational approaches: discovery of GTPase HRas as prospective drug target for 1,3-diazine scaffolds. *BMC Chem*, **13**, 96.

95. Brand,S.E., Scharlau,M., Geren,L., Hendrix,M., Parson,C., Elmendorf,T., Neel,E., Pianalto,K., Silva-Nash,J., Durham,B., *et al.* (2022) Accelerated Evolution of Cytochrome c in Higher Primates, and Regulation of the Reaction between Cytochrome c and Cytochrome Oxidase by Phosphorylation. *Cells*, **11**, 4014.

96. Sirawaraporn,W., Sathitkul,T., Sirawaraporn,R., Yuthavong,Y. and Santi,D.V. (1997) Antifolate-resistant mutants of Plasmodium falciparum dihydrofolate reductase. *Proc Natl Acad Sci U S A*, **94**, 1124–1129.
